# Detecting Introgression in Shallow Phylogenies: How Minor Molecular Clock Deviations Lead to Major Inference Errors

**DOI:** 10.1101/2025.03.25.645378

**Authors:** Xiao-Xu Pang, Jianquan Liu, Da-Yong Zhang

**Affiliations:** College of Ecology, Lanzhou University, Lanzhou 730000, China; Ministry of Education Key Laboratory for Biodiversity Science and Ecological Engineering, College of Life Sciences, Beijing Normal University, Beijing 100875, China

**Keywords:** *D*-statistic, HyDe, false positive, introgression, phylogenomics, rate variation, shallow phylogenies

## Abstract

Recent theoretical and algorithmic advances in introgression detection, coupled with the growing availability of genome-scale data, have highlighted the widespread occurrence of interspecific gene flow across the tree of life. However, current methods largely depend on the molecular clock assumption—a questionable premise given empirical evidence of substitution rate variation across lineages. While such rate heterogeneity is known to compromise gene flow detection among divergent lineages, its impact on closely related taxa at shallow evolutionary timescales remains poorly understood, likely because these taxa are often assumed to adhere to a molecular clock. To address this gap, we combine theoretical analyses and simulations to evaluate the robustness of widely used site-pattern methods (*D*-statistic and HyDe) to rate variation across phylogenetic timescales. Our results demonstrate that both methods exhibit high sensitivity to even minor deviations from the molecular clock at shallow timescales, complementing previous findings at deeper scales. Specifically, in young phylogenies (with an age of 3×10^5^ generations) with small population sizes, weak (17% difference) and moderate (33% difference) rate variation can inflate false-positive rates up to 35% and 100%, respectively, when using site-pattern counts from a 500Mb genome. Employing a more distant outgroup intensifies these spurious signals. Our study demonstrates that summary tests for introgression are pervasively vulnerable to minor rate variations and underscores the critical need for advanced methodologies to disentangle genuine introgression from false signals generated by rate heterogeneity.

## Introduction

Mounting genomic evidence indicates that interspecific gene flow—the transfer of genetic material between species through hybridization—occurs far more extensively than previously recognized across the tree of life (Mallet et al. 2016; Suarez-Gonzalez et al. 2018; Taylor and Larson 2019; Edelman and Mallet 2021). As Edelman and Mallet (2021, p.271) observed, “phylogenies with no evidence of gene flow are beginning to seem like the exception rather than the rule.” Testing for gene flow (or introgression) has become a routine component of phylogenetic analyses, enabling researchers to evaluate hybridization’s role in a group’s species diversification and to guide the selection between tree-based and network-based evolutionary frameworks (Blair and Ané 2020; Haque and Kubatko 2024).

Over the past few years, various introgression detection approaches have been developed within a rigorous hypothesis-testing framework (Jiao et al. 2021; Hibbins and Hahn 2022). These methods typically require a predefined species tree, and can be classified into two main categories based on the type of information utilized. The first category relies on gene-tree topologies or parsimony-informative site patterns, detecting gene flow through significant asymmetries between discordant gene trees or between discordant site patterns, as exemplified by the widely-used *D*-statistic (Green et al. 2010), HyDe (Blischak et al. 2018) and MSCquartets (Rhodes et al. 2021). The second utilizes gene-tree branch lengths as a source of information, examining whether the branch-length distributions are influenced by introgression events. Examples include *D*_*3*_ (Hahn and Hibbins 2019), QuIBL (Edelman et al. 2019), and the full-likelihood tests that make full use of both topological and branch length information in gene trees (Zhu and Yang 2012; Ji et al. 2023). Particularly, site pattern-based methods operate directly on the whole genome sequence alignment data, thus avoiding potential errors introduced by preprocessing steps such as loci partitioning and gene-tree estimation.

Despite significant advances in introgression testing methods, each approach relies, to varying degrees, on various theoretical assumptions, and violations of these assumptions can lead to erroneous conclusions. One such assumption—the molecular clock or rate constancy among lineages, which is either implicitly or explicitly built into many widely used methods—has recently garnered considerable attention (Blair and Ané 2020; Frankel and Ané 2023; Koppetsch et al. 2024). For instance, *D*-statistic is based on the infinite-site mutation model that each site undergoes only one mutation (Green et al. 2010). Under this assumption, *ABBA* and *BABA* site patterns can only arise from a single substitution event along the internal branches of discordant gene trees. Gene flow is the only possible mechanism disrupting the *ABBA*-*BABA* balance, which is otherwise maintained by incomplete lineage sorting (ILS) (Pamilo and Nei 1988; Degnan 2013). However, these discordant site patterns may also result from homoplasies, where identical alleles arise independently in different lineages. When rate variation exists between sister lineages, homoplasies can create *ABBA*-*BABA* asymmetry, mimicking introgression. Such false positives in the *D*-statistic have been confirmed in recent simulation studies by Frankel and Ané (2023) and Koppetsch et al. (2024), which examined cases involving deep divergences where sister lineages have been separated for over a million generations. However, it remains unclear whether this same caveat applies to shallow phylogenies, where only moderate deviations from the clock assumption are possible at best. Additionally, a significant limitation of these studies is their reliance solely on simulations, without theoretical analysis to support their findings.

Substitution rates are shaped by various factors such as generation time, effective population size, metabolic rate, and longevity (Bromham 2009; Thomas et al. 2010; Lanfear et al. 2014). Since some of these factors can vary considerably over short timescales, substitution rates may differ significantly even among closely related species, as evidenced by numerous empirical studies (Bromham et al. 2013; Dann et al. 2017; Saclier et al. 2018; Bergeron et al. 2023). To better understand the prevalence and magnitude of rate variation within shallow phylogenies, we employed the relative rate test (Graur and Li 2000) to quantify rate difference between a pair of lineages and applied it to published datasets from a random selection of 6 genera (see Supplementary Note 1), spanning both plant and animal taxa (Leduc-Robert and Maddison 2018; Wu et al. 2018; Karimi et al. 2020; Sarver et al. 2021; Liu et al. 2022; Gardner et al. 2023). Notably, intra-generic species frequently exhibit rate disparities of 10%–30%, with some species pairs in *Habronattus* and *Malus* even exceeding 50% (Fig. 1). Thus, rate variation among closely related species is both widespread and appreciable, underscoring the need to account for such variation in evolutionary analyses. Moreover, the *D*-statistic was originally designed for testing hybridization between humans and Neanderthals, which diverged about 20,000 generations ago (Green et al. 2010), and remains predominantly applied to groups with comparable evolutionary scales (i.e., within tens of thousands to a few millions of generations). Given the widespread use of *D*-statistic and other site pattern-based methods in shallow phylogenies, it is critical to assess and quantify their reliability in the presence of rate variation across lineages.

**Figure 1.**
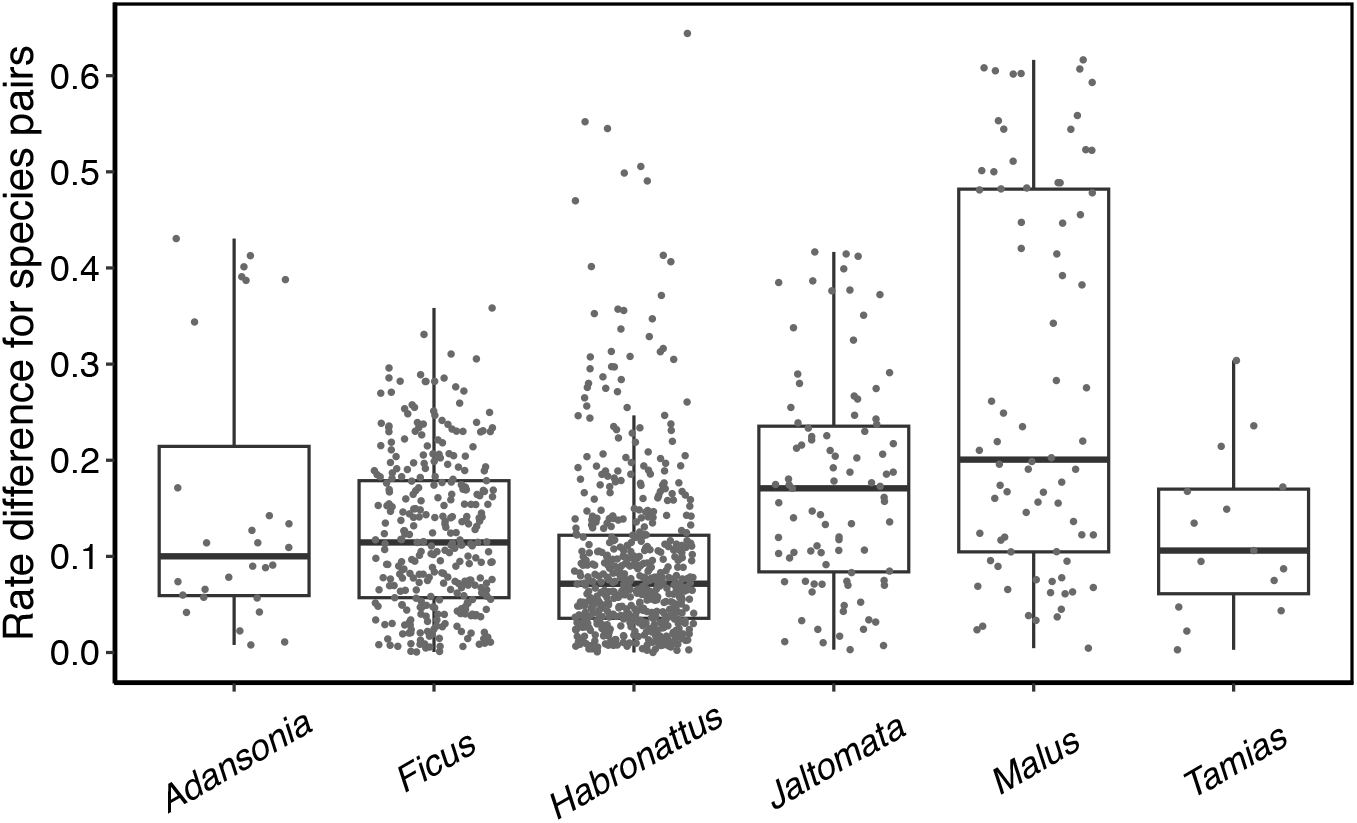
Evolutionary rate differences among species pairs across six genera, calculated using the relative rate test (see Graur and Li 2000, Supplementary Note 1). Each point represents the rate difference between a pair of intra-generic species. Here, the rate difference is defined as the difference in lengths between two branches divided by the length of the longer branch.

Our study aims to assess and quantify the robustness of site pattern-based *D*-statistic and HyDe methods against lineage-specific rate variation at shallow phylogenetic scales. We began with a mathematical analysis to derive the expected *D*-values under varying degrees of rate variation between sister lineages, along with other key parameters such as phylogenetic age, effective population size, and outgroup distance. We then simulated a range of scenarios spanning phylogenetic depths of 10^4^ to 10^6^ generations to examine the behaviors of *D*-statistic and HyDe in the presence of rate variation. Our results reveal the widespread conditions under which rate variation generates false-positive signals of introgression in popular summary tests, calling into question the validity of numerous reported gene-flow events, which may, in fact, be artifacts of substitution rate heterogeneity.

### Theory

We begin with an introduction to the basic principles of the site pattern-based methods *D*-statistic (Green et al. 2010) and HyDe (Blischak et al. 2018). In the four-taxon asymmetric species tree ***S*** with the topology of (((*P1, P2*), *P3*), *O*), when introgression is absent, 2 discordant unrooted gene trees caused by ILS, *p1p3*|*p2o* and *p2p3*|*p1o*, occur with equal probabilities but at lower frequencies than *p1p2*|*p3o* (Pamilo and Nei 1988; Degnan 2013). Building on this property, *D*-statistic and HyDe assume an infinite-site model (i.e., no multiple hits) and use parsimony-informative site patterns—*ABBA, BABA*, and *BBAA* (with *A* representing the reference state from the outgroup listed last)—as proxies for 3 gene-tree topologies. Introgression is detected by identifying deviations from the expected symmetry between *ABBA* and *BABA* counts. Specifically, the *D*-statistic, calculated as:

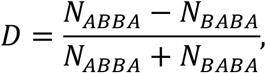

detects introgression when the *D*-value significantly deviates from 0. When *D* > 0, *D*-statistic suggests introgression between *P2* and *P3*, leading to an excess of *ABBA* over *BABA*; conversely, when *D* < 0, it points to introgression between *P1* and *P3*. HyDe employs a different statistic (refer to Kubatko and Chifman 2019 for further details) but fundamentally tests whether the 2 least frequent site patterns occur at comparable frequencies. A significant result is interpreted as indicative of a hybrid speciation event, where the 2 parental species are more distantly related to each other than to the hybrid species. Thus, the 2 ingroup lineages that share alleles in the smallest number of site patterns will be identified as putative parents, with the remaining lineage considered the hybrid (Kubatko and Chifman 2019). It should be noted that, while ghost introgression can also yield significant *D*-values (Tricou et al. 2022; Pang and Zhang 2024), here we do not incorporate ghost lineages into consideration for the sake of conceptual simplicity.

In our previous study (Pang and Zhang 2024), the frequencies of 3 parsimony-informative site patterns are derived under the infinite-site model. Here, we relax this assumption to allow for homoplasies and further incorporate rate variation between sister lineages. We use the multispecies coalescent with introgression (MSci; Flouri et al. 2020) model to consider an episodic introgression event from *P*1 to *P*3 within the species tree ***S***, with the introgression proportion denoted by *γ* (Fig. 2a). Speciation and introgression times, *τ*_*i*_ = *T*_*i*_μ, and population sizes, *θ* = 4*N*μ, are measured in terms of the expected number of mutations per site, where *T*_*i*_ is the time in generations, *N* is the number of diploid individuals, assumed constant across lineages, and μ represents the mutation rate per site per generation. To account for rate variation between a pair of sister lineages, we introduce a relative rate parameter *λ* for lineage *P1*, scaling its branch length to *λτ*_1_ and population size to *λθ* accordingly.

**Figure 2.**
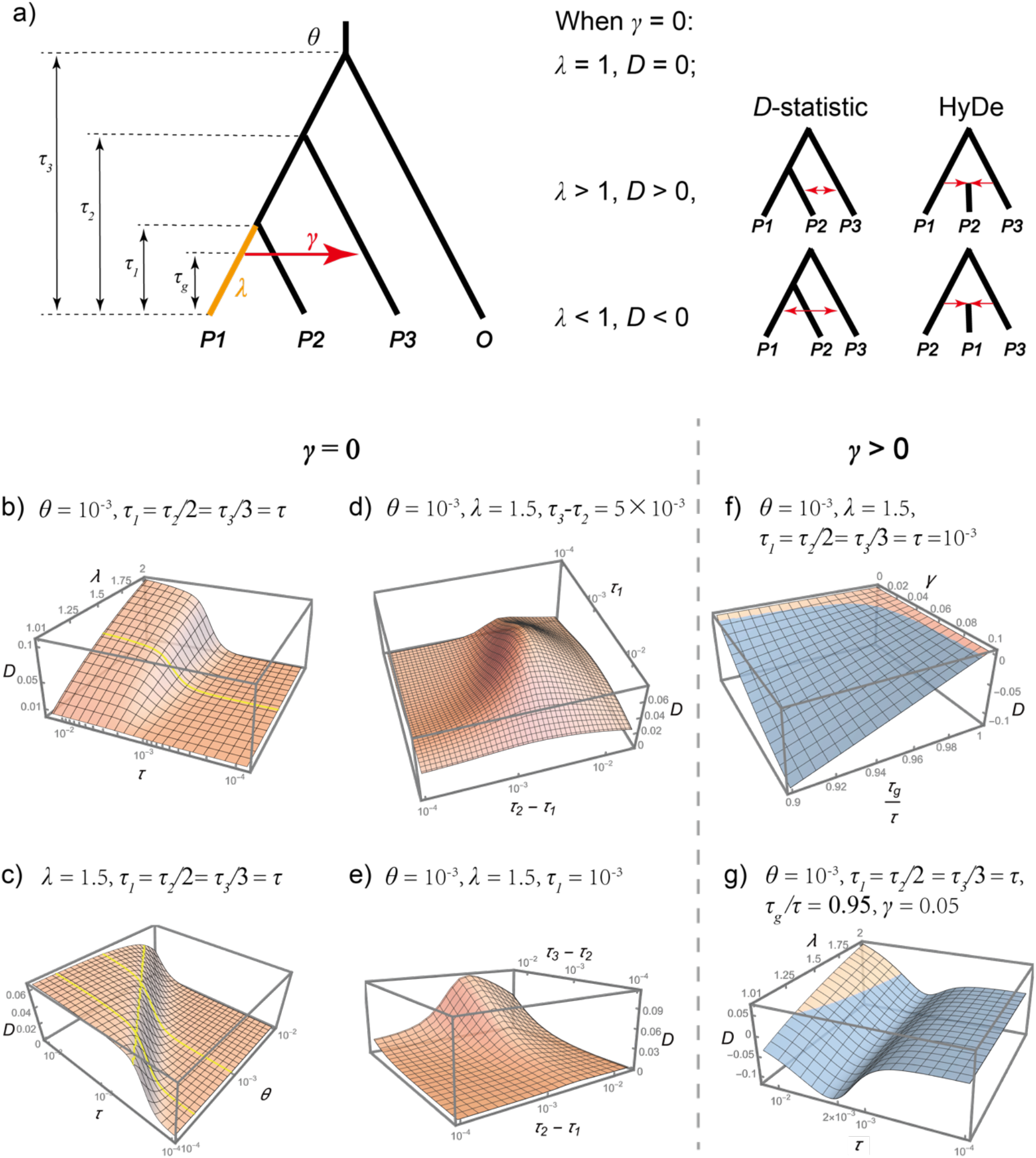
Impact of rate variation on introgression testing. a) Left panel: A four-taxon asymmetric species tree with an outflow introgression (*P1*→*P3*) and rate variation between sister lineages. The introgression probability is denoted as *γ*. Divergence time *τ*_*i*_ {*i* = 1, 2, 3}, introgression time *τ*_*g*_, and the ancestral population size *θ* are measured in terms of the expected number of mutations per site. The lineage *P1*, highlighted in orange, exhibits a distinct substitution rate relative to other lineages, with *λ* representing its relative rate. Right panel: Expected behaviors of *D*-statistic and HyDe under different conditions of *λ* in the absence of introgression (*γ* = 0). b-e) Plots illustrating the effect of various parameters on *D*-values in the absence of introgression (*γ* = 0). Axis for *τ* and *θ* are log-scaled. The yellow curve in panel (b) marks *λ* = 1.5. Yellow curves in panel (c) correspond to *θ* = 2 × 10^−4^, *θ* = 10^−3^, and *D* = 0.05. f-g) Plots illustrating the effect of various parameters on *D*-values in the presence of introgression (*γ* > 0). Orange and blue surfaces in 3D plots indicate parameter regions where *D* > 0 and *D* < 0, respectively.

Gene tree distributions are calculated under the MSci model, and sequence evolution along gene trees is modeled using the Jukes–Cantor (JC69) substitution model (Jukes and Cantor 1969). The frequencies of 3 site patterns per nucleotide site are derived by traversing all possible gene-tree histories and are provided in Supplementary Note 2.

#### Case 1: γ = 0 (No introgression)

We first consider the case where *γ* = 0. The expected frequency difference between *ABBA* and *BABA* is given by:

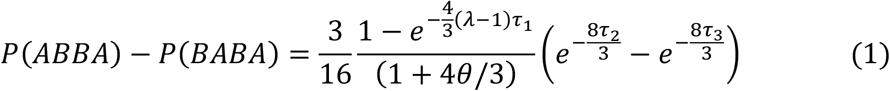

where *τ*_3_ > *τ*_2_ > *τ*_1_, *τ*_*i*_, *λ* ∈ (0, ∞). It follows that the relative magnitudes of *ABBA* and *BABA* counts are determined by the parameter *λ*. Additionally, the inequality *P*(*BBAA*) > *P*(*BABA*) necessarily holds, as demonstrated in Supplementary Note 3. From these observations, we can draw the following conclusions:

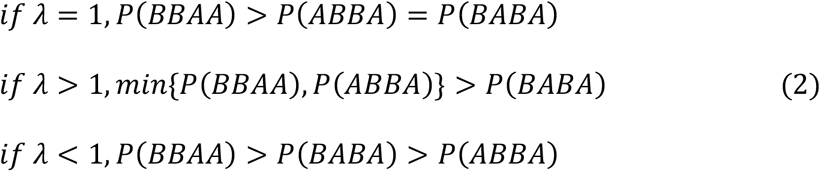

When *λ* = 1 (i.e., no rate variation), the *ABBA-BABA* balance is maintained (*D* = 0) even in the presence of homoplasies. However, when *λ* ≠ 1, this balance is disrupted. Specifically, when *λ* > 1, indicating an accelerated rate for *P1* lineage, *D* > 0, *D*-statistic suggests introgression between lineages *P2* and *P3*, while HyDe infers *P1* and *P3* as the parental lineages—those sharing alleles in the fewest site pattern *BABA*—and *P2* as the hybrid lineage (Fig. 2a). Conversely, when *λ* < 1, reflecting a slower rate of *P1*, there is *D* < 0, *D*-statistic suggests introgression between *P1* and *P3*, while HyDe would classify *P2* and *P3* as the parental lineages— those sharing the derived state in the fewest site pattern *ABBA*—and *P2* as the hybrid lineage (Fig. 2a).

Quantitatively, the absolute difference between *ABBA* and *BABA* primarily depends on the terms 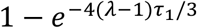 and 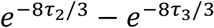 in Equation (1), which reflect the length differences between sister lineages *P1* and *P2*, and between *P3* and the outgroup *O*, respectively. Obviously, greater length differences augment the extent of *ABBA*-*BABA* asymmetry. Consider the extreme cases where *P1* and *O* exhibit long branches while *P2* and *P3* have negligible ones (i.e., *τ*_1_, *τ*_2_ → 0, and *λτ*_1_, *τ*_3_ → ∞), the majority of homoplasies occur along the *P1* and *O* lineages to generate *ABBA* sites. Consequently, *D*-value approaches 1 if *θ* ≪ 1 (detailed proof in Supplementary Note 4). Conversely, if the *P1–P2* or *P3–O* pair has equal branch lengths (i.e., *λ* = 1 or *τ*_2_ = *τ*_3_), *D* equals 0.

Next, we analyze the effects of various parameters on the *D*-value. First, we plot the *D* against *λ* and *τ*, assuming *τ*_1_ = *τ*_2_/2 = *τ*_3_/3 = *τ* and fixing *θ* = 10^−3^ (Fig. 2b). The results indicate that *D* generally increases with *λ* and *τ*, in line with the expectation that greater rate variation and deeper phylogeny amplify spurious introgression signals. Specifically, *D* shows an approximately linear increase with *λ*, while follows an S-shaped curve with respect to *τ*, featuring a sharp rise around *τ* = 10^−3^ and leveling off near *τ* = 3 × 10^−3^. For example, with *λ* = 1.5 (Fig. 2b), increasing *τ* from 10^−3^ to 3 × 10^−3^ results in a pronounced rise in *D* from 0.006 to 0.055; further increasing *τ* to 10^−2^ only causes a slight increase in *D*, reaching a value of 0.063.

Then, we plot the *D* against *τ* and *θ*, fixing *λ* = 1.5 (Fig. 2c). The results reveal a strong interaction between *θ* and *τ* in determining *D*-values, where smaller *θ* values necessitate lower phylogenetic depth (*τ*) thresholds for a high *D*-value. For instance, when *θ* = 10^−3^, a *τ* value of 2.4 × 10^−3^ is required to reach *D* = 0.05, whereas for *θ* = 2 × 10^−4^, only 5.8 × 10^−4^ of *τ* is needed (Fig. 2c). This pattern arises because larger *θ* amplifies ILS, which contributes equally to *ABBA* and *BABA*, thereby diluting their difference caused by homoplasies.

Finally, we evaluate the effects of individual branch lengths, specifically *τ*_1_, *τ*_2_−*τ*_1_ and *τ*_3_−*τ*_2_, while fixing *θ* = 10^−3^ and *λ* = 1.5. When *D* is plotted against *τ*_1_ and *τ*_2_−*τ*_1_, with *τ*_3_−*τ*_2_ = 5 × 10^−3^ fixed, it exhibits a unimodal pattern for both variables, initially increasing and then decreasing (Fig. 2d). The initial rises with *τ*_1_ and *τ*_2_−*τ*_1_ are driven by an accumulation of homoplasy-induced sites and a reduction in ILS-induced sites, respectively (Fig. S3). The subsequent decline in *D*, though counterintuitive, can be explained by Equation (1): Increasing both variables elongates the branch lengths of *P3* and *O* (i.e., *τ*_2_ and *τ*_3_), thereby reducing the value of the term 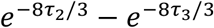 and diminishing the *ABBA-BABA* difference, despite an increased occurrence of homoplasies. When *D* is plotted against the outgroup distance *τ*_3_−*τ*_2_ with *τ*_1_ = 10^−3^ fixed, *D* increases with larger values of *τ*_3_−*τ*_2_ (Fig. 2e). This effect is driven by increasing branch length disparities between *P3* and the outgroup *O*, which intensify homoplasy-induced asymmetry.

#### Case 2: γ > 0 (introgression present)

We next examine the cases where *γ* > 0, indicating the presence of gene flow. First, we plot the *D* against *γ* and *τ*_*g*_/*τ*, with *τ* = *τ*_1_ = *τ*_2_/2 = *τ*_3_/3 = 10^−3^, *θ* = 10^−3^, and *λ* = 1.5 fixed (Fig. 2f). The results show that both the degree and timing of introgression, *γ* and *τ*_*g*_/*τ*, have a significant impact on *D*-values. More intense and recent introgression strengthen introgression signals, making them less susceptible to rate variation. Only in a limited parameter space, where *γ* < 0.03 or *τ*_*g*_/*τ* > 0.98, does rate variation result in *D* close to 0 or even positive (as indicated by the orange region of Fig. 2f).

We then fix *τ*_*g*_/*τ* = 0.95 and *γ* = 0.05, and plot *D* against *λ* and *τ* to explore the effect of phylogenetic age (Fig. 2g). When *λ* = 1, as expected, *D* < 0. Interestingly, the absolute value of *D* follows a non-monotonic pattern, initially increasing and then decreasing as *τ* increases. This implies that introgression signals are strongest at moderate phylogenetic depths. Further increases in *λ* lead to a rise in *D*, and for larger *τ*, the sign of *D* even reverses from negative to positive (Fig. 2g).

### Simulations

To assess the robustness of introgression testing methods to rate variation among lineages, we simulated 8 different scenarios, each with distinct focus points. Simulations were conducted under the species tree ***S***, either with or without introgression (Figs. 3a, 7a). Time parameters (*TS, TI, TO*, and *TG*) were measured in generations, and *λ* represents the relative rate of a specific lineage. To compare the quantitative effects of rate acceleration and deceleration, we defined the rate difference as the difference in lengths between two branches divided by the length of the longer branch. Thus, both *λ* = *x* (*x* > 1) and *λ* = 1/*x* correspond to a rate difference of 1 − 1/*x*. We examined 4 relative rate pairs: 0.2 vs. 5, 0.5 vs. 2, 0.67 vs. 1.5, and 0.83 vs. 1.2, with *λ* = 1 serving as the baseline. Detailed parameter configurations for each scenario are provided in Table 1.

**Table 1.**
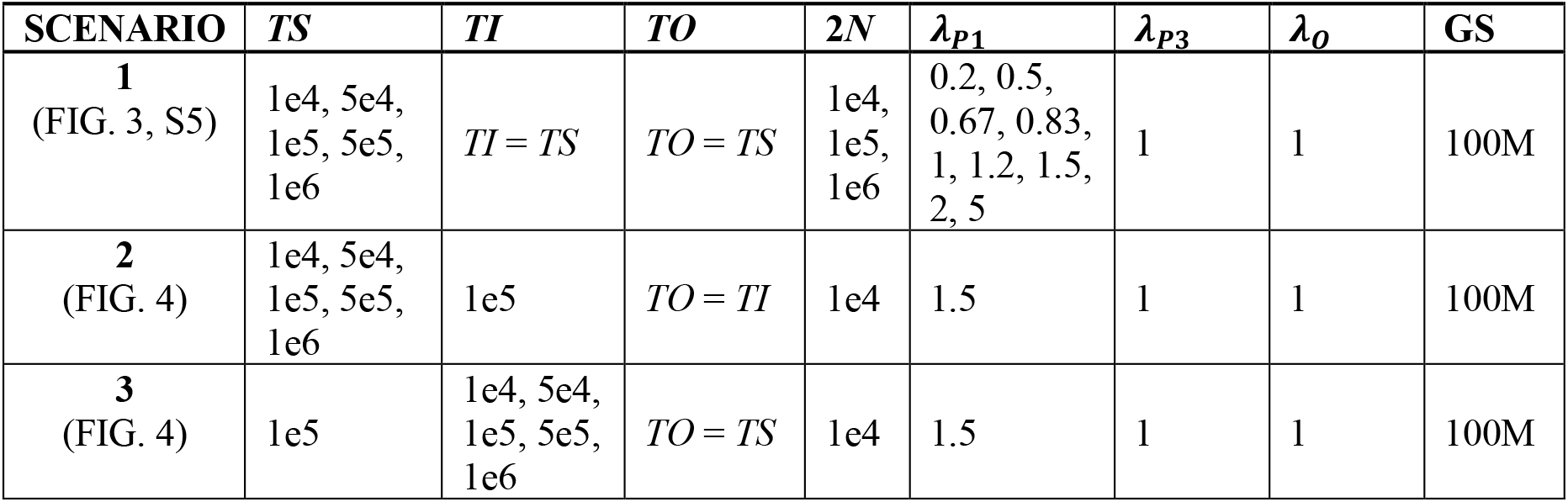
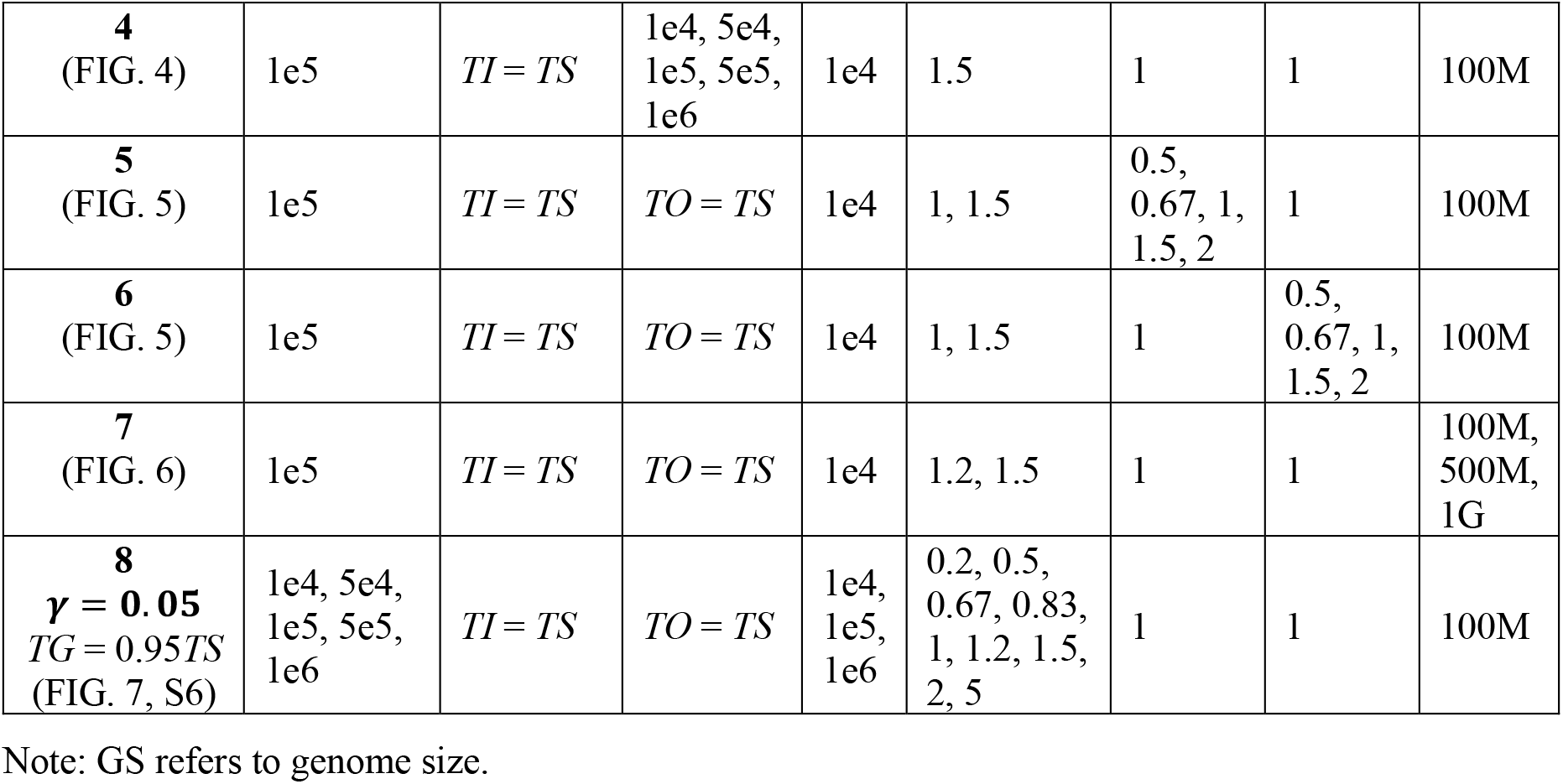
The simulation scenarios and parameter settings.

| SCENARIO | $TS$ | $TI$ | $TO$ | $2N$ | $\lambda_{P1}$ | $\lambda_{P3}$ | $\lambda_O$ | GS |
| --- | --- | --- | --- | --- | --- | --- | --- | --- |
| **1**<br>(FIG. 3, S5) | 1e4, 5e4,<br>1e5, 5e5,<br>1e6 | $TI = TS$ | $TO = TS$ | 1e4,<br>1e5,<br>1e6 | 0.2, 0.5,<br>0.67, 0.83,<br>1, 1.2, 1.5,<br>2, 5 | 1 | 1 | 100M |
| **2**<br>(FIG. 4) | 1e4, 5e4,<br>1e5, 5e5,<br>1e6 | 1e5 | $TO = TI$ | 1e4 | 1.5 | 1 | 1 | 100M |
| **3**<br>(FIG. 4) | 1e5 | 1e4, 5e4,<br>1e5, 5e5,<br>1e6 | $TO = TS$ | 1e4 | 1.5 | 1 | 1 | 100M |
| **4**<br>(FIG. 4) | 1e5 | $TI = TS$ | 1e4, 5e4,<br>1e5, 5e5,<br>1e6 | 1e4 | 1.5 | 1 | 1 | 100M |
| **5**<br>(FIG. 5) | 1e5 | $TI = TS$ | $TO = TS$ | 1e4 | 1, 1.5 | 0.5,<br>0.67, 1,<br>1.5, 2 | 1 | 100M |
| **6**<br>(FIG. 5) | 1e5 | $TI = TS$ | $TO = TS$ | 1e4 | 1, 1.5 | 1 | 0.5,<br>0.67, 1,<br>1.5, 2 | 100M |
| **7**<br>(FIG. 6) | 1e5 | $TI = TS$ | $TO = TS$ | 1e4 | 1.2, 1.5 | 1 | 1 | 100M,<br>500M,<br>1G |
| **8**<br>$\gamma = 0.05$<br>$TG = 0.95TS$<br>(FIG. 7, S6) | 1e4, 5e4,<br>1e5, 5e5,<br>1e6 | $TI = TS$ | $TO = TS$ | 1e4,<br>1e5,<br>1e6 | 0.2, 0.5,<br>0.67, 0.83,<br>1, 1.2, 1.5,<br>2, 5 | 1 | 1 | 100M |

Note: GS refers to genome size.

**Figure 3.**
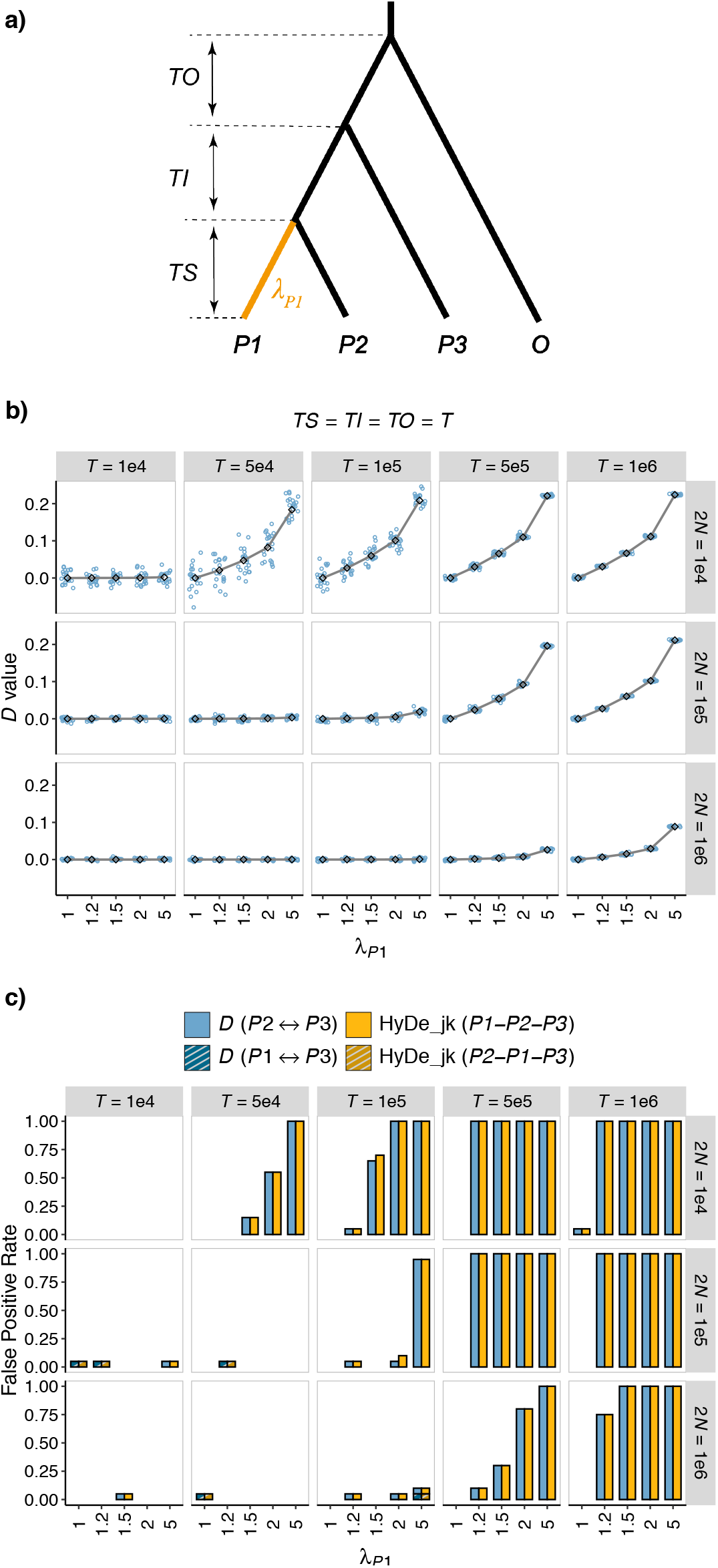
Interactive effects of phylogenetic depth (*T*), population size (2*N*), and rate variation on summary tests for introgression. a) Simulated demographic histories. Branch lengths (*TS, TI*, and *TO*) are measured in generations. *λ*_*P*1_ represents the relative rate of the lineage *P1*. b-c). Results of introgression tests under the condition where *TS* = *TI* = *TO* = *T*. The values on the strips at the top and right of each plot indicate the phylogenetic depth (*T*) and population size (2*N*), respectively. *λ*_*P*1_ is labeled on the x-axis. b) *D*-values: Colored points represent *D* estimates, with solid lines indicating the theoretical expected *D*. c) False-positive rate of *D*-statistic and HyDe_jk.

#### Note: GS refers to genome size

For each scenario, we first used fastsimcoal v2.8 (Excoffier et al. 2013) to simulate gene trees, with single sample per lineage. Since fastsimcoal2 does not natively support evolutionary rate variation among lineages, we accounted for this by scaling branch lengths by *λ* through controlled sampling times of samples. For example, if the evolutionary rate of the *P1* lineage was accelerated (*λ*_*P*1_ > 1), all divergence times were shifted into the past by (*λ*_*p*1_ − 1) * *TS*, with *P1* sampled at the present while all other extant lineages sampled at (*λ*_*P*1_ − 1) * *TS* generations ago. We then simulated 100-bp-long sequences from each gene tree using Seq-Gen software (Rambaut and Grass 1997) under the JC69 model and a mutation rate μ = 2 × 10^−8^ per site per generation (“-s 2e-8”). The per-gene sequences were concatenated to generate genome sizes of 100 Mb, unless otherwise specified. Each parameter combination was simulated with 20 replications.

*D*-statistic and HyDe methods were applied to each dataset. HyDe v.0.4.3 (Blischak et al. 2018) was run using the ‘run_hyde.py’ script, with *O* designated as the outgroup. Since HyDe assumes site independence, we implemented a block-jackknife approach via a custom Python script to account for correlations introduced by shared genealogies, referring to this modified version as “HyDe_jk.” For *D*-statistic, we calculated *D*-values based on site pattern counts provided by HyDe and estimated the standard error *SE(D)* using the block-jackknife method. The significance of introgression was assessed via *z* = *D*/*SE*(*D*), with a *p*-value threshold of 0.01.

## Results

### Effect of phylogeny age and population size

In this section, we simulated the scenarios without introgression (Fig. 3a, parameters: Scenario 1 in Table 1). Assuming branch lengths *TS = TI = TO = T*, we varied the parameter *T* and 2*N* to investigate how phylogenetic depth interacts with population size to influence the occurrence of false positives caused by rate variation. The results are summarized in Figure 3 and S5.

In the absence of rate variation (*λ*_01_ = 1), *D*-values, as expected, remained close to 0, and the FP rates of *D*-statistic and HyDe were low across all parameter combinations. When the evolutionary rate of the *P1* lineage accelerated (*λ*_*P*1_ > 1), *D*-values increased and became positive (Fig. 3b). Under these conditions, *D*-statistic detected significant signals of introgression between the *P2* and *P3* lineages in many parameter combinations, and HyDe misidentified the *P2* lineage as the hybrid (Fig. 3c). On the other hand, when the evolutionary rate of the *P1* lineage slowed (*λ*_*P*1_< 1), *D*-values turned negative, suggesting introgression between the *P1* and *P3* lineages. In this case, HyDe misidentified the *P2* lineage as the hybrid (Fig. S5). These results align with our theoretical predictions illustrated in Figure 2a. Notably, we observed that for the same degree of rate variation (i.e., *λ*_*P*1_ = *k* versus *λ*_*P*1_ = 1/*k*), a decrease in the evolutionary rate of *P1* leads to FP rates similar to those observed when the rate is accelerated, though slightly lower.

We found that *D*-values deviated further from 0, and FP rates increased with greater phylogenetic depth (*T*) and smaller population sizes (2*N*). For example, when 2*N* = 10^4^ and *T* = 10^5^, a 50% increase of substitution rate (*λ*_*P*1_ = 1.5, corresponding to 33% rate difference) resulted in a mean *D*-value of 0.063 (range: 0.0279 to 0.1097) and approximately 70% FP rate (Figs. 3b-c). In contrast, with a larger population size of 2*N* = 10^5^ and *λ*_*P*1_ = 1.5, the mean *D*-value dropped to around 0 (range: −0.01304 to 0.0117), and the FP rate became negligible for *T* ≤ 10^5^.

### Effects of divergence time of sister lineages, internal branch length and distance of outgroup

We further investigated the effects of individual branch lengths *TS, TI*, and *TO*. In these analyses, we considered scenarios (Scenarios 2-4 in Table 1) with a rate variation *λ*_*P*1_ = 1.5 and a population size of 2*N* = 10^4^. We initialized *TS* = *TI* = *TO* = 10^5^ generations, and independently varied each branch length across a range of 10^4^ to 10^6^ generations to assess its specific effects on introgression testing outcomes. The results are displayed in Figure 4.

**Figure 4.**
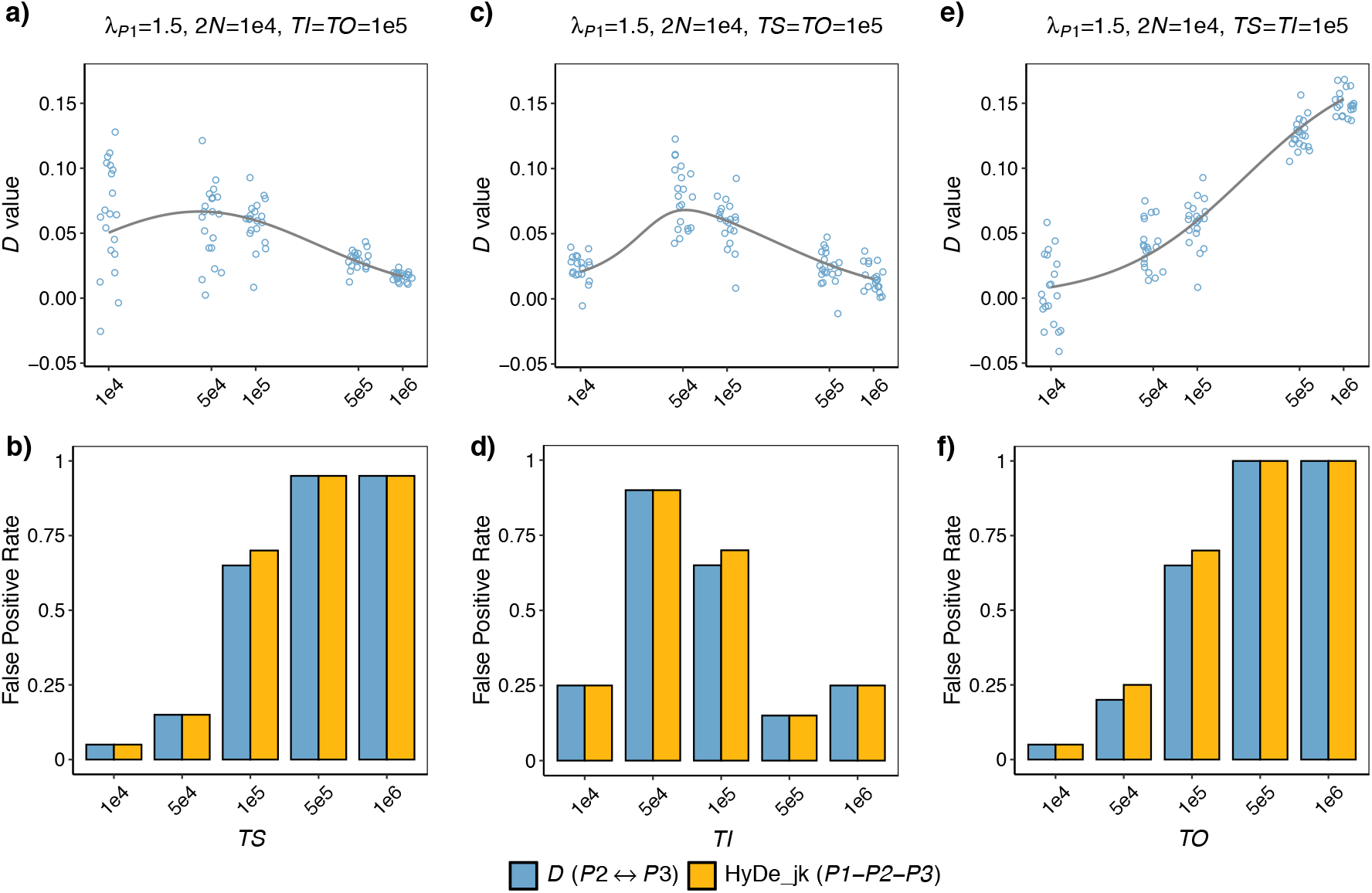
Interactive effects of individual branch lengths (*TS, TI*, and *TO*) and rate variation on summary tests for introgression. The demographic scenario corresponds to Figure 3a, with *λ*_*P*1_ = 1.5 and 2*N* = 10^1^. Default branch lengths are set to *TS* = *TI* = *TO* = 10^5^ generations. a-b) *D*-values and false-positive rates with varying *TS*. Colored points in panel (a) represent *D*-value estimates, with solid lines indicating the theoretical expected *D*-values. c-d) Results with varying *TI*. e-f) Results with varying *TO*.

First, we examined the effect of the divergence time of sister lineages *TS*. The *D*-values initially increased with *TS* but showed a decline when *TS* > 5×10^4^ (Fig. 4a). This later decline trend was also observed in a study by Koppetsch et al. (2024) in the context of deep divergence of sister lineages (>10 million generations). Interestingly, the FP rates exhibited a different trend, steadily increasing as *TS* increased (Fig. 4b). FP rates were approximately 15% at *TS* = 5×10^4^ and reached nearly 100% when *TS* = 10^6^, even though the mean *D*-value was obviously higher at *TS* = 5×10^4^ (0.059) compared to *TS* = 10^6^ (0.017). This discrepancy may arise because larger *TS* values contribute more homoplasy-induced *ABBA* and *BABA* sites, reducing sample variance, as reflected by the lower variance of *D*-values across replicates. This, in turn, results in a higher z-score, despite the slight decline in *D*-values. Next, for the internal branch length *TI*, both *D*-values and FP rates followed a non-monotonic trend: initially rising with *TI* and then decreasing, peaking at *TI* = 5×10^4^ with a mean *D*-value of 0.078 and FP rate of approximately 75% (Fig. 4c-d). Finally, for the outgroup distance *TO, D*-values gradually deviated from 0 with increasing *TO*, reaching a striking value of ∼0.15 at *TO* = 10^2^ (Fig. 4e). FP rates followed a similar trend, rising from nearly 0% at *TO* = 10^4^, to ∼70% at *TO* = 10^1^, and reaching 100% at *TO* ≥ 5 × 10^1^ (Fig. 4f). This suggests that a larger distance between the outgroup and the ingroups exacerbates the likelihood of false positives due to rate variation.

### Effects of P3- or O-mutation rate variation

This section extends the analysis to the effects of rate variation in *P3* and the outgroup *O* on introgression testing. We considered the following scenarios (Scenarios 5-6 in Table 1) with branch lengths of *TS* = *TI* = *TO* =10^5^ generations, and a population size of 2*N* = 10^4^. The relative rates of *P3* and *O, λ*_*p*3_ and *λ*_3_, were varied under conditions with and without rate variation between sister lineages. The corresponding results are presented in Figure 5.

**Figure 5.**
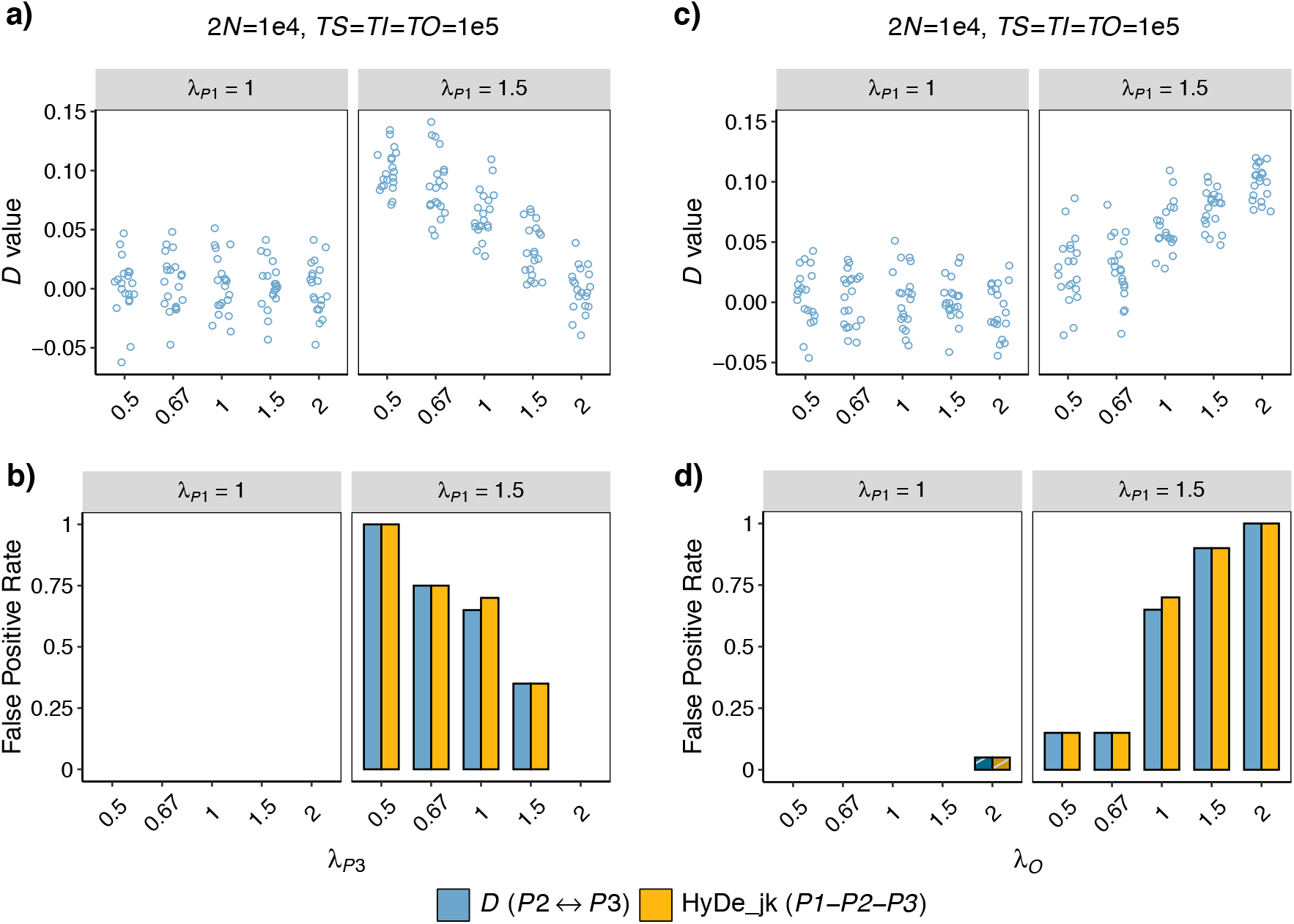
Effects of *P3*- and *O*-rate variation on summary tests for introgression. The simulated demographic scenario corresponds to Figure 3a, with additional rate variation in lineages *P3* and *O*, where *λ*_*P*3_ and *λ*_*O*_ denote their relative rates. Branch lengths are fixed at *TS* = *TI* = *TO* = 10^5^ generations, and 2*N* = 10^4^. a-b) *D*-values and false-positive rates with varying *λ*_4−_. The top strips of each plot indicate conditions for *λ*_*P*1_ = 1 and *λ*_*P*1_= 1.5, corresponding to scenarios without and with rate variation between sister lineages, respectively. Colored points in panel (a) represent *D*-value estimates. c-d) Results with varying *λ*_*O*_.

In the absence of rate variation between sister lineages (i.e., *λ*_43_ = 1), the estimated *D*-values remained consistently around 0 regardless of changes in the rates of *P3* or *O*. In this case, all methods performed well, with the rejection rates of the *D*-statistic and HyDe-jk at 1% and 1.5%, respectively (Fig. 5).

When rate variation was present among sister lineages (i.e., *λ*_*P*1_ = 1.5), reducing the *P3*-mutation rate led to a noticeable increase in both *D*-values and FP rates (Fig. 5a-b). For instance, a 50% reduction in the *P3*-mutation rate (i.e., *λ*_*P*3_= 0.5) increased the mean *D*-value from 0.063 to 0.01 and the FP rate from ∼70% to 100%. Conversely, doubling *P3*-rate (i.e., *λ*_*P*3_= 2) reduced the mean *D*-value to nearly 0 and eliminated false positives. The effects of rate variation in the outgroup *O* followed an opposite trend (Figs. 5c-d). Halving the *O*-rate (*λ*_*O*_ = 0.5) decreased the mean *D*-value to 0.027 and the FP rate to 15%, while doubling the *O*-rate (*λ*_*O*_ = 2) increased the mean *D*-value to 0.01 and the FP rate to 100%.

### Effects of Genome Size (or Amount of Data)

In statistical testing, larger sample sizes generally enhance statistical power. Given the wide variation in genome sizes across animal and plants, ranging from 12 Mb to 27.6 Gb, with a median of 517 Mb and a mean of 1.21 Gb (Kress et al. 2022), our initial genome size of 100 Mb may be too small. Here, in Scenarios 7 (Table 1) with branch lengths of *TS* = *TI* = *TO =T* ranging from 10^4^ to 10^5^ generations, rate variation of *λ*_*P*1_ = 1.2 or 1.5, and a population size of 2*N* = 10^4^, we simulated larger genome sizes of 500 Mb and 1 Gb to explore the effects of genome size on the rate variation-induced FP rates.

The results show that, as expected, the estimated *D*-values across replicates become less scattered as genome size increased, converging toward the theoretically expected value (indicated by the grey lines). Meanwhile, the FP rates of both methods rise significantly with genome size. Under weak rate variation of *λ*_*P*1_ = 1.2 and *T* = 10^5^ generations, expanding the dataset size from 100 Mb to 500 Mb and 1 Gb elevated the FP rate from 5% to approximately 35% and 60%, respectively. Under stronger rate variation (*λ*_*P*1_ = 1.5), FP rates reached ∼100% at just 500 Mb. These results highlight that larger dataset sizes make introgression testing methods more susceptible to rate variation.

### False negatives and positives caused by rate variation in introgression scenarios

In this part, we simulated introgression scenarios (Fig. 7a, Scenarios 8 in Table 1). We assumed branch lengths *TS = TI = TO = T*, and considered an episodic outflow event from *P1* to *P3* at 0.95*T*, with an introgression proportion of 0.05. This analysis focuses on the role of phylogenetic depth (*T*) and population size (2*N*) on introgression testing. Figures 7 and S6 summarizes the corresponding results.

When rate variation was absent (i.e., *λ*_*p*1_ = 1), *D*-values were negative as expected. *D*-statistic successfully detected introgression between *P1* and *P3* under many parameter combinations (Fig. 7b-c). HyDe, however, confused the direction of gene flow, misidentifying *P1* as the hybrid lineage, an issue having already been noted in previous studies (Pang and Zhang 2024). The degree to which *D*-values deviated from 0 depended on both phylogenetic age and population size. For phylogenetic age *T*, the relationship between *D*-values and *T* showed a U-shaped trend (Fig. 7b). Especially in cases with a small population size of 2*N* = 10^4^, the mean *D*-values sharply decreased from -0.0078 to -0.19 as *T* increased from 10^4^ to 5×10^4^, then gradually increased to -0.03 at *T* = 10^6^. The power of tests rose from 5% at *T* = 10^4^ to 100% for *T* ≥ 5×10^4^ (Fig. 7c). Regarding population size, both the absolute *D*-values and the test power decreased as 2*N* increased, indicating that true introgression signals are more readily detected in groups with small ancestral population sizes (Figs. 7b-c).

When *P1*’s mutation rate accelerated (i.e., *λ*_*P*1_ > 1), *D*-values gradually shifted from negative to 0 and even became positive as *λ*_*P*1_ increased (Fig. 7b). This trend resulted in a decreased test power, and in some cases, the detection of spurious introgression events (Fig. 7c). This negative effect of rate variation was particularly pronounced in cases with older phylogenies and small population sizes. For example, when *T* = 10^6^ and 2*N* = 10^4^, a 20% increase in the evolutionary rate of *P1* (*λ*_*P*1_ = 1.2) led to a failure to detect introgression signals (10% power), while a 50% increase (*λ*_*P*1_ = 1.5) resulted in incorrect inference of *P2↔P3* gene flow using the *D*-statistic (100% FP rate) (Fig. 7c). In contrast, when the evolutionary rate of *P1* was slower (i.e., *λ*_*P*1_ < 1), rate variation had a beneficial effect, enhancing the *ABBA-BABA* asymmetry and improving the power to detect *P1↔P3* introgression (Fig. S6).

## Discussion

The widespread use of summary tests for introgression has spurred numerous evaluation studies on their performance, including sensitivity analyses for false negatives (Zheng and Janke 2018; Kong and Kubatko 2021; Bjorner et al. 2024) and robustness assessments against false positives (Tricou et al. 2022; Frankel and Ané 2023; Koppetsch et al. 2024; Pang and Zhang 2024). Recently, attention has been drawn to the commonly assumed clock model in these methods, with simulation studies revealing that rate variation can introduce false-positive signals in deeply divergent groups (Frankel and Ané 2023; Koppetsch et al. 2024). Here, we shift the focus to shallow phylogenies, where rate variation remains widespread and appreciable (Fig. 1) and where introgression testing is most commonly applied. We systematically evaluate the sensitivity of site pattern-based methods, *D*-statistic and HyDe, to rate variation under various phylogenetic and demographic conditions.

Our theoretical analysis confirms that rate variation between sister lineages can create asymmetric *ABBA* and *BABA* counts, even in the absence of introgression (Fig. 2a). Specifically, the slower-evolving sister lineage has a higher probability of sharing alleles with *P3*. This may appear counterintuitive and contradicts the conclusions of Frankel and Ané (2023; Figure 8), who suggest that the faster-evolving sister lineage and *P3* are more likely to undergo homoplasies and share alleles. The discrepancy arises because Frankel and Ané (2023) ignored mutations along the outgroup *O*, which necessarily occur more frequently than those along *P3*. As a consequence, rate variation-induced *ABBA-BABA* imbalance is misinterpreted as a signal of gene flow, with the slower-evolving sister lineage mistakenly identified as a participant in an introgression event by *D*-statistic and as a hybrid lineage by HyDe. Furthermore, in cases under introgression, rate variation can alter the magnitude or even the sign of the *D*-value, potentially leading to both false negatives and false positives (Fig. 7).

Phylogeny age interacts with population size to influence the occurrence of false positives caused by rate variation. Our results support the intuition that introgression tests performed for older groups are more vulnerable to rate variation (Fig. 3). Nevertheless, phylogenetic age is not the sole determinant, because population size also plays a key role. Specifically, small population sizes (in ancestral lineages) can render shallow phylogenies sensitive to rate variation, producing high *D*-values and false-positive signals. For example, in young groups with a phylogeny age of 3×10^5^ generations (corresponding to *T* = 10^5^ in Fig. 3 and S5) and a small population size of 2*N* = 10^4^, even a weak 17% rate variation between sister lineages (i.e., *λ* = 1.2) results in FP rates of 35% for 500 Mb datasets, rising to 60% for 1 Gb datasets (Fig. 6b). A moderate 33% rate difference (i.e., *λ* = 0.67 or 1.5) produces *D*-values exceeding 0.05, comparable to the 0.047 *D*-value reported in the Neanderthal-modern human introgression study by Green et al. (2010). Under this condition, FP rates surpass 50% in a 100 Mb dataset (Fig. 3 and S5) and reach 100% when the dataset size increases to 500 Mb (Fig. 6). Moreover, the risk of false positives is expected to worsen when using population-level data with multiple individuals per lineage. Interestingly, a previous work by Zheng and Janke (2018) also revealed the pivotal role of population size in the power of the *D*-statistic when introgression is present, showing a significant negative correlation between the two. In conclusion, both false and true introgression signals are stronger in groups with smaller population sizes. Larger population sizes, in contrast, produce higher ILS-driven *ABBA* and *BABA* counts increasing the prevalence of discordant gene tree topologies and elongating their internal branches, which may obscure asymmetric signals caused by either rate variation or actual introgression.

**Figure 6.**
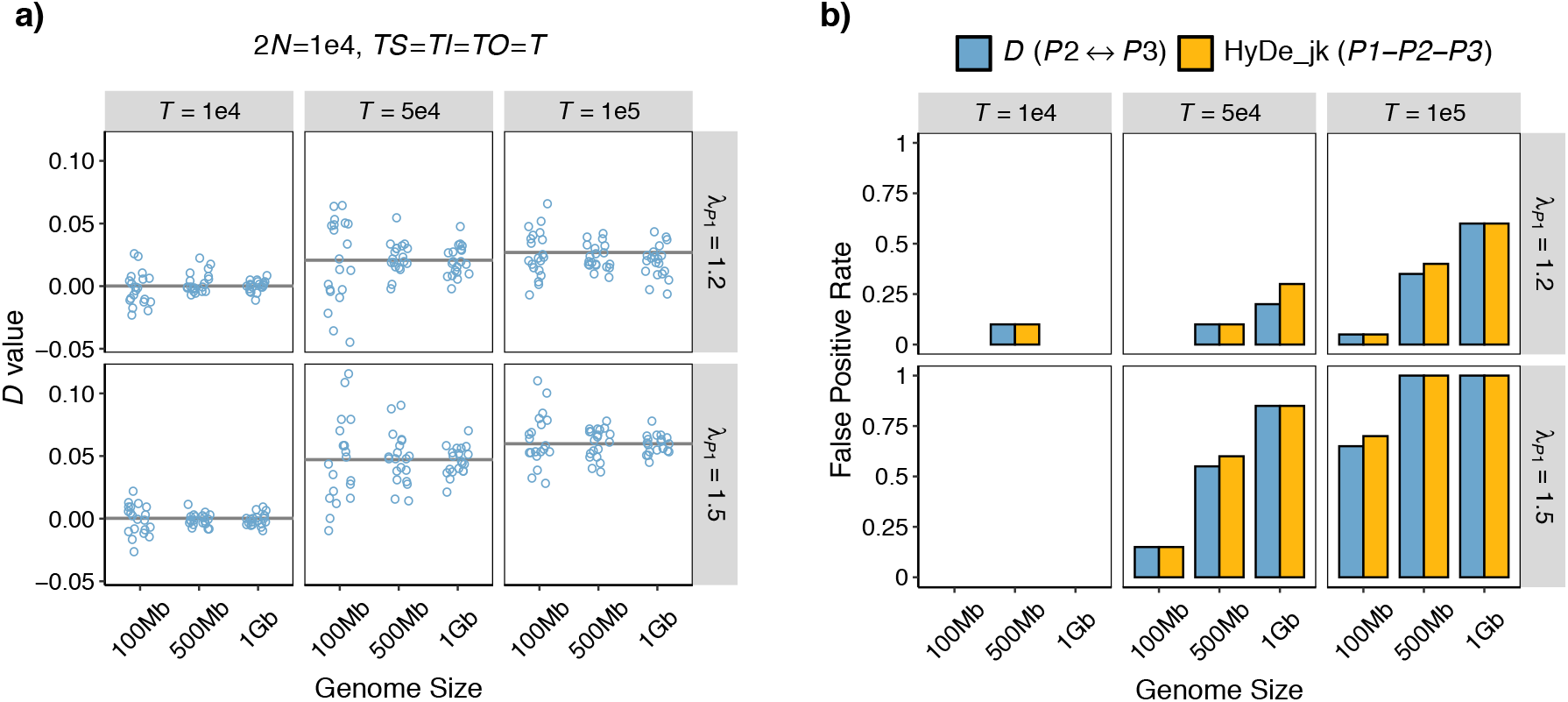
Effects of dataset size on summary tests for introgression. The simulated demographic scenario corresponds to Figure 3a, Branch lengths are fixed at *TS* = *TI* = *TO* = *T*, and 2*N* = 10^1^. The values on the strips at the top and right of each plot indicate the phylogenetic depth (*T*) and the extent of rate variation (*λ*_*P*1_), respectively. Genome size is labeled on the x-axis. a) *D*-values: Colored points represent *D* estimates, with solid lines indicating the theoretical expected *D*. b) False-positive rate of *D*-statistic and HyDe_jk.

**Figure 7.**
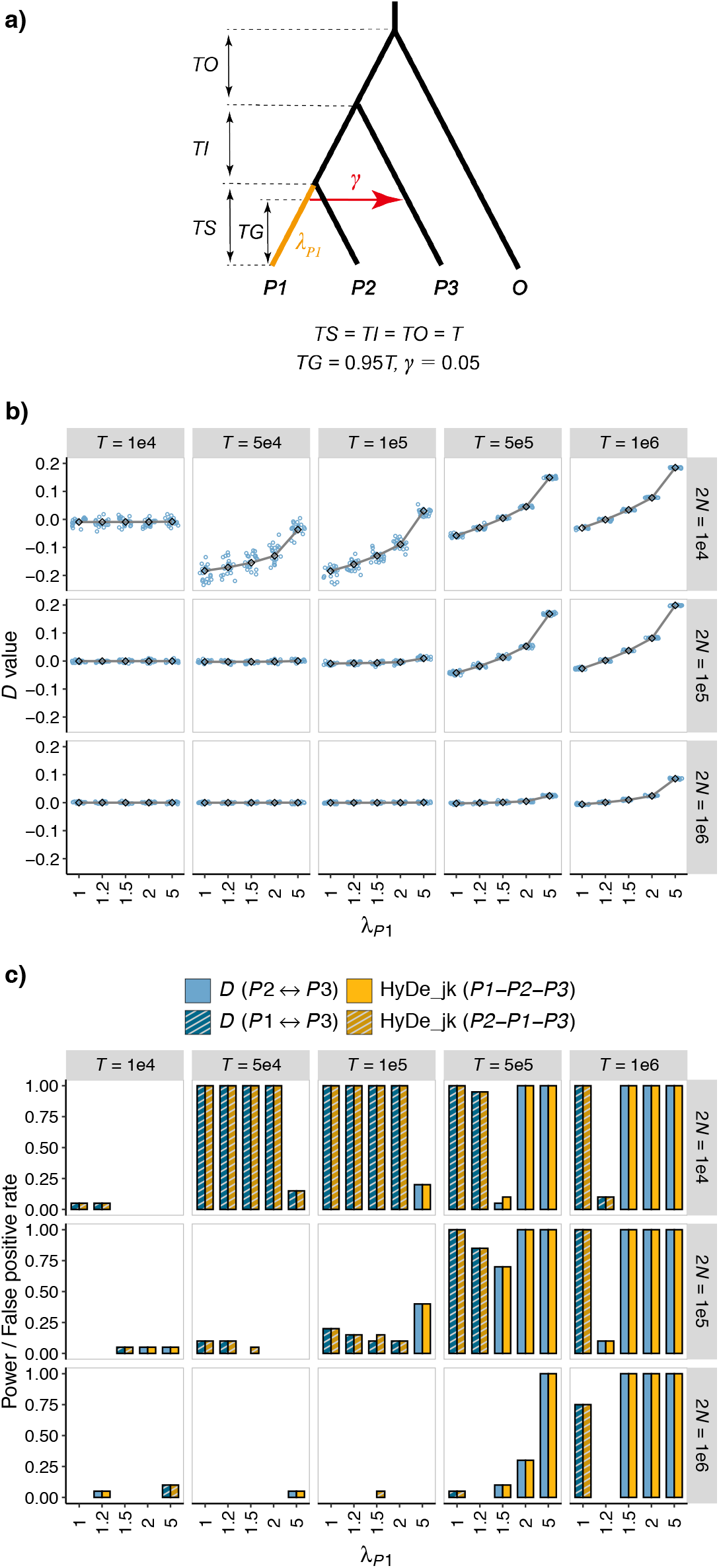
Effect of rate variation on summary tests, in presence of introgression. a) Simulated demographic histories. Branch lengths are measured in generations, with *TS* = *TI* = *TO* = *T*. An outflow introgression (*P1*→*P3*) occurs at 0.95*T* with a proportion *γ* of 0.05. *λ*_*P*1_ represents the relative rate of lineage *P1*. b-c). Results of introgression tests. The values on the strips at the top and right of each plot indicate the phylogenetic depth (*T*) and population size (2*N*), respectively. *λ*_*P*1_ is labeled on the x-axis. b) *D*-values: Colored points represent *D*-value estimates, with solid lines indicating the theoretical expected *D*-values. c) False-positive rate of *D*-statistic and HyDe_jk.

Outgroup distance has a large impact on the incidence of false positives due to rate heterogeneity. In empirical studies, the outgroup is typically selected to ensure sufficient genetic divergence from the ingroups to mitigate the risk of introgression between them (Green et al. 2010). Zheng and Janke (2018) justified this practice, suggesting that the power of the *D*-statistic seems unaffected by outgroup distance. However, their simulations did not account for rate variation among lineages. Our study shows that a more distant outgroup can largely increase the likelihood of false positives arising from rate variation (Figs. 2d and 4e-f). For instance, in the scenarios discussed earlier (*TS* = *TI* = *TO* = 10^5^ generations and *λ* = 1.5), increasing the outgroup distance *TO* to 10^6^ generations leads to a striking increase in the *D*-value to around 0.15, alongside a FP rate of 100% based on a 100 Mb dataset (Fig. 4e-f). This occurs because a larger branch difference between *P3* and the outgroup *O* amplifies *ABBA-BABA* asymmetry resulting from homoplasies. Due to the same underlying mechanism, a reduced evolutionary rate of *P3* or an accelerated rate in the outgroup *O* can also produce a similar effect of exacerbating FP rates (Fig. 5). Our findings highlight the challenges in selecting an appropriate outgroup, necessitating careful consideration of the trade-offs between distant and closely related outgroups. We lean toward recommending the selection of a closely related outgroup, as this can yield higher-quality data due to fewer mapping and alignment errors, while also reducing false positives caused by rate variation. On the other hand, researchers should also be mindful of the potential for introgression involving the outgroup when interpreting significant results.

Our study did not evaluate methods that rely on gene-tree branch lengths or those that depend exclusively on gene-tree topologies. Intuitively, the former is more susceptible to deviations from the clock assumption, while the latter is likely more robust (Frankel and Ané 2023; Koppetsch et al. 2024). This expectation stems from the fact that rate variation does not alter the coalescent trajectories of alleles, but rather only affects mutation accumulation along these coalescent branches. When rate variation causes significant fluctuations in branch lengths, deviating substantially from the expected distribution under ILS, spurious introgression signals may be introduced to account for these discrepancies. For example, Frankel and Ané (2023) found that *D*_*3*_ is particularly vulnerable to rate variation between two sister lineages. We anticipate that full-likelihood tests would also exhibit high sensitivity to rate heterogeneity, which warrants further investigation to confirm. As for topology-based methods, their performance depends heavily on the accuracy of gene-tree topology inference. These methods may perform well for deep phylogenies, but are less suitable for shallow phylogenies where the sparse phylogenetic signal per locus impedes accurate topological reconstruction. For example, loci exhibiting only a single parsimony-informative site, if arising from homoplasy, inevitably produce inaccurate topologies due to long-branch attraction.

### Implications for Practice

In empirical systems, researchers may wonder under what conditions introgression testing is most susceptible to rate variation and how to differentiate it from genuine introgression signals. For a given group of closely related taxa, we recommend first quantifying the extent of rate variation using the relative rate test, as applied to the 6 genera in the Introduction section of this study (Fig. 1). Particular caution is warranted when species pairs exhibit more than 20% rate differences, especially if they have small ancestral population sizes, as these factors together significantly elevate false-positive risks. Additionally, we advise conducting simulations tailored to the group’s specific characteristics, including the extent of rate variation, divergence times, population size, and genome size. Such simulations can offer a more nuanced view of rate variation’s impact on the group in question.

Several strategies may help researchers distinguish between patterns driven by rate variation and those caused by genuine introgression. A key feature of rate variation-driven signals is that the sister lineage with the shorter branch length tend to have a higher probability of sharing alleles with *P3*. Therefore, if a test detects introgression between the faster-evolving sister lineage and *P3*, this strongly suggests that the signal reflects genuine introgression rather than an artifact of rate variation. Furthermore, Bayesian model comparison offers a rigorous framework for evaluating the evidence for gene flow in light of violating the clock assumption. Species tree methods, which are well-established and flexible, can accommodate variable assumptions about rate variation across lineages. A practical approach involves comparing two models under a full-likelihood framework: (1) a species tree model that accounts for rate heterogeneity (e.g., a relaxed clock model) and (2) an introgression model that explicitly incorporates gene flow. If the introgression model demonstrates a significantly higher marginal likelihood than the rate-heterogeneity model, this outcome provides statistical support for the presence of gene flow. However, this strategy is computationally demanding may be infeasible for large genome-scale datasets.

Koppetsch et al. (2024) recently developed a user-friendly “*ABBA*-site clustering” test to distinguish between spurious and genuine introgression signals, based on the distribution of *D*-informative sites across the genome. The test capitalizes on the fact that introgression typically leaves behind haplotypes with clusters of linked “*ABBA*” sites that reflect introgression history, while sites arising from homoplasies are expected to be randomly distributed. By identifying *ABBA*-clustering patterns along chromosomes, the test can effectively detect introgression in deep phylogenies. However, mutation hotspots or mapping biases can also generate clustered “*ABBA*” sites. To address this, Koppetsch et al. (2024) developed a more “robust” version of the test, but it exhibited high false-negative rates even under strong introgression. Additionally, its performance in shallow phylogenies remains uncertain, where mutations are rarer and introgression-driven clusters may be difficult to detect.

Given these limitations, there is a pressing need for more refined and powerful methods to distinguish between signals caused by rate variation and those arising from genuine introgression. Until such methods are developed, empirical researchers should adopt a conservative approach by considering rate variation as a default explanation when interpreting significant results from introgression tests.

## Author Contributions

Xiao-Xu Pang and Da-Yong Zhang jointly conceived and designed the study, and drafted the manuscript. Xiao-Xu Pang performed theoretical analyses and simulations. Jianquan Liu revised the manuscript, and all authors approved the final version.

## Acknowledgements

We would like to thank Yang Yang and Wei-Ning Bai for helpful discussions. This work was supported by China Postdoctoral Science Foundation (GZB20240286) and Beijing Advanced Innovation Program for Land Surface Processes.

## Data Availability Statement

The simulation results generated in this study, as well as the scripts and files necessary to run the analyses, are available via Data Dryad:

## Reference

Bergeron LA, Besenbacher S, Zheng J, Li P, Bertelsen MF, Quintard B, Hoffman JI, Li Z, St. Leger J, Shao C, et al. 2023. Evolution of the germline mutation rate across vertebrates. Nature 615:285–291.

Bjorner M, Molloy EK, Dewey CN, Solis-Lemus C. 2024. Detectability of varied hybridization scenarios using genome-scale hybrid detection methods. Bulletin of the Society of Systematic Biologists 3(1).

Blair C, Ané C. 2020. Phylogenetic trees and networks can serve as powerful and complementary approaches for analysis of genomic data. Syst. Biol. 69:593–601.

Blischak PD, Chifman J, Wolfe AD, Kubatko LS. 2018. HyDe: A python package for genome-scale hybridization detection. Syst. Biol. 67:821–829.

Bromham L. 2009. Why do species vary in their rate of molecular evolution? Biol. Lett. 5:401–404.

Bromham L, Cowman PF, Lanfear R. 2013. Parasitic plants have increased rates of molecular evolution across all three genomes. BMC Evol. Biol. 13:126.

Dann M, Bellot S, Schepella S, Schaefer H, Tellier A. 2017. Mutation rates in seeds and seed-banking influence substitution rates across the angiosperm phylogeny. bioRxiv 10.1101/156398.

Degnan JH. 2013. Anomalous unrooted gene trees. Syst. Biol. 62:574–590.

Edelman NB, Frandsen PB, Miyagi M, Clavijo B, Davey J, Dikow RB, García-Accinelli G, Van Belleghem SM, Patterson N, Neafsey DE. 2019. Genomic architecture and introgression shape a butterfly radiation. Science 366:594–599.

Edelman NB, Mallet J. 2021. Prevalence and adaptive impact of introgression. Annu. Rev. Genet. 55:265–283.

Excoffier L, Dupanloup I, Huerta-Sanchez E, Sousa VC, Foll M. 2013. Robust demographic inference from genomic and SNP data. PLoS Genet. 9:e1003905.

Flouri T, Jiao X, Rannala B, Yang Z. 2020. A Bayesian implementation of the multispecies coalescent model with introgression for phylogenomic analysis. Mol. Biol. Evol. 37:1211–1223.

Frankel LE, Ané C. 2023. Summary tests of introgression are highly sensitive to rate variation across lineages. Syst. Biol. 72:1357–1369.

Gardner EM, Bruun-Lund S, Niissalo M, Chantarasuwan B, Clement WL, Geri C, Harrison RD, Hipp AL, Holvoet M, Khew G. 2023. Echoes of ancient introgression punctuate stable genomic lineages in the evolution of figs. Proc. Natl. Acad. Sci. U.S.A. 120:e2222035120.

Graur D, Li W-HL. 2000. Fundamentals of molecular evolution. 12th ed. Sunderland, MA: Sinauer.

Green RE, Krause J, Briggs AW, Maricic T, Stenzel U, Kircher M, Patterson N, Li H, Zhai W, Fritz MH-Y. 2010. A draft sequence of the Neandertal genome. Science 328:710–722.

Hahn MW, Hibbins MS. 2019. A three-sample test for introgression. Mol. Biol. Evol. 36:2878–2882.

Haque MR, Kubatko L. 2024. A global test of hybrid ancestry from genome-scale data. Stat. Appl. Genet. Mol. Biol. 23:20220061.

Hibbins MS, Hahn MW. 2022. Phylogenomic approaches to detecting and characterizing introgression. Genetics 220:iyab173.

Ji J, Jackson DJ, Leaché AD, Yang Z. 2023. Power of Bayesian and heuristic tests to detect cross-species introgression with reference to gene flow in the Tamias quadrivittatus group of north American chipmunks. Syst. Biol. 72:446–465.

Jiao X, Flouri T, Yang Z. 2021. Multispecies coalescent and its applications to infer species phylogenies and cross-species gene flow. Natl. Sci. Rev. 8:nwab127.

Jukes TH, Cantor CR. 1969. Evolution of protein molecules. Mammalian protein metabolism. New York: Academic Press. p. 21–132.

Karimi N, Grover CE, Gallagher JP, Wendel JF, Ané C, Baum DA. 2020. Reticulate evolution helps explain apparent homoplasy in floral biology and pollination in baobabs (Adansonia; Bombacoideae; Malvaceae). Syst. Biol. 69:462–478.

Kong S, Kubatko LS. 2021. Comparative performance of popular methods for hybrid detection using genomic data. Syst. Biol. 70:891–907.

Koppetsch T, Malinsky M, Matschiner M. 2024. Towards reliable detection of introgression in the presence of among-species rate variation. Syst. Biol. 73:769–788.

Kress WJ, Soltis DE, Kersey PJ, Wegrzyn JL, Leebens-Mack JH, Gostel MR, Liu X, Soltis PS. 2022. Green plant genomes: What we know in an era of rapidly expanding opportunities. Proc. Natl. Acad. Sci. U.S.A. 119:e2115640118.

Kubatko LS, Chifman J. 2019. An invariants-based method for efficient identification of hybrid species from large-scale genomic data. BMC Evol. Biol. 19:1–13.

Lanfear R, Kokko H, Eyre-Walker A. 2014. Population size and the rate of evolution. Trends Ecol. Evol. 29:33–41.

Leduc-Robert G, Maddison WP. 2018. Phylogeny with introgression in Habronattus jumping spiders (Araneae: Salticidae). BMC Evol. Biol. 18:1–23.

Liu BB, Ren C, Kwak M, Hodel RG, Xu C, He J, Zhou WB, Huang CH, Ma H, Qian GZ. 2022. Phylogenomic conflict analyses in the apple genus Malus sl reveal widespread hybridization and allopolyploidy driving diversification, with insights into the complex biogeographic history in the Northern Hemisphere. J. Integr. Plant Biol. 64:1020–1043.

Mallet J, Besansky N, Hahn MW. 2016. How reticulated are species? Bioessays 38:140–149.

Pamilo P, Nei M. 1988. Relationships between gene trees and species trees. Mol. Biol. Evol. 5:568–583.

Pang X-X, Zhang D-Y. 2024. Detection of ghost introgression requires exploiting topological and branch length information. Syst. Biol. 73: 207–222.

Rambaut A, Grass NC. 1997. Seq-Gen: An application for the Monte Carlo simulation of DNA sequence evolution along phylogenetic trees. Bioinformatics 13:235–238.

Rhodes JA, Baños H, Mitchell JD, Allman ES. 2021. MSCquartets 1.0: Quartet methods for species trees and networks under the multispecies coalescent model in R. Bioinformatics 37:1766–1768.

Saclier N, François CM, Konecny-Dupré L, Lartillot N, Guéguen L, Duret L, Malard F, Douady CJ, Lefébure T. 2018. Life history traits impact the nuclear rate of substitution but not the mitochondrial rate in isopods. Mol. Biol. Evol. 35:2900–2912.

Sarver BA, Herrera ND, Sneddon D, Hunter SS, Settles ML, Kronenberg Z, Demboski JR, Good JM, Sullivan J. 2021. Diversification, introgression, and rampant cytonuclear discordance in rocky mountains chipmunks (Sciuridae: Tamias). Syst. Biol. 70:908–921.

Suarez-Gonzalez A, Lexer C, Cronk QC. 2018. Adaptive introgression: A plant perspective. Biol. Lett. 14:20170688.

Taylor SA, Larson EL. 2019. Insights from genomes into the evolutionary importance and prevalence of hybridization in nature. Nat. Ecol. Evol. 3:170–177.

Thomas JA, Welch JJ, Lanfear R, Bromham L. 2010. A generation time effect on the rate of molecular evolution in invertebrates. Mol. Biol. Evol. 27:1173–1180.

Tricou T, Tannier E, de Vienne DM. 2022. Ghost lineages highly influence the interpretation of introgression tests. Syst. Biol. 71:1147–1158.

Wu M, Kostyun JL, Hahn MW, Moyle LC. 2018. Dissecting the basis of novel trait evolution in a radiation with widespread phylogenetic discordance. Mol. Ecol. 27:3301–3316.

Zheng Y, Janke A. 2018. Gene flow analysis method, the D-statistic, is robust in a wide parameter space. BMC Bioinform. 19:10.

Zhu T, Yang Z. 2012. Maximum likelihood implementation of an isolation-with-migration model with three species for testing speciation with gene flow. Mol. Biol. Evol. 29:3131–3142.

