## supplemental materials for "Detecting Introgression in Shallow Phylogenies: How Minor Molecular Clock Deviations Lead to Major Inference Errors"

**Supplementary Note 1**

We employed the relative rate test (Graur and Li 2000) to quantify the extent of rate variation between species pairs. Specifically, we used an outgroup, *O*, as reference. For each lineage pair within the ingroups, *A* and *B*, we calculated the branch lengths of *AI* and *BI* for each locus, based on genetic distances of the sequence pairs $K_{AB}$, $K_{AO}$, and $K_{BO}$, as described in Figure S1. The genetic distances were calculated using the *dist.dna* function from the **ape** R package, with the JC69 model and the option pairwise.deletion = TRUE. Then we computed the mean value of $K_{AI}$ and $K_{BI}$ across all loci, with their relative difference representing the rate difference of the species pair *A* and *B*.

It should be noted that the branch lengths of *AI* and *BI* for each locus include a segment corresponding to the common ancestor of species *A* and *B*, where the evolutionary rate is shared. Therefore, the relative rate test method tends to underestimate rate difference.

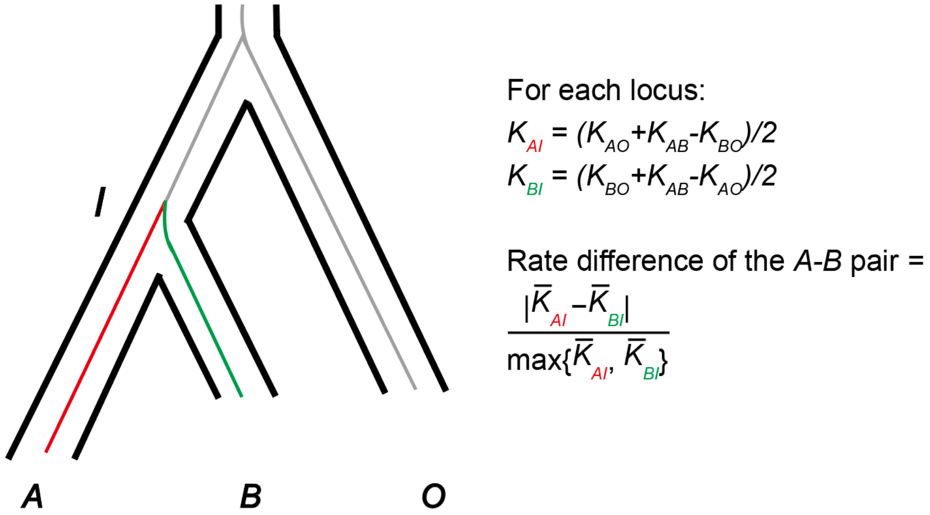


Figure S1. Calculation of rate difference between species pairs (Graur and Li 2000).

Table S1. Details of the Six Genera Datasets

| Genera | Number of species | Number of Loci | Literatures |
|---|---|---|---|
| *Adansonia* | 8 | 372 | (Karimi et al. 2020) |
| *Ficus* | 26 | 1477 | (Gardner et al. 2023) |
| *Habronattus* | 34 | 1877 | (Leduc-Robert and Maddison 2018) |
| *Jaltomata* | 14 | 1000 | (Wu et al. 2018) |
| *Malus* | 13 | 620 | (Liu et al. 2022) |
| *Tamias* | 6 | 1060 | (Sarver et al. 2021) |

### Supplementary Note 2

For the MSci model presented in Figure S1 (right), since there is only one descendant from the hybrid node, the coalescent process can be decomposed into two parental trees with probabilities $\gamma$ and $1-\gamma$, respectively (Meng and Kubatko 2009; Zhu and Degnan 2017) (Fig. S1). The probabilities of site patterns for a four-taxon tree under conditions of rate variation have been derived by Richards and Kubatko (2022). The site-pattern frequencies of the MSci model are expressed as a linear combination of those from the two parental trees (i.e., *S* and *I*), weighted by their respective probabilities:

$$P(site_i|MSci,\lambda) = (1-\gamma)P(site_i|S,\lambda) + \gamma P(site_i|I,\lambda).$$

Let $k = e^{-(4/3)(\lambda-1)\tau_1}, k' = e^{-(4/3)(1-\lambda)\tau_g}, a = 4\theta/3$, the frequencies of three parsimony-informative site pattern per site for outflow introgression are given in Table S2.

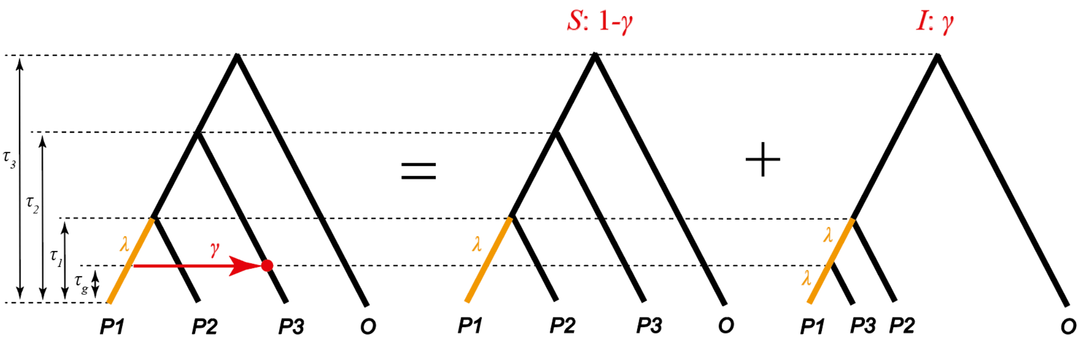


Figure S2. Decomposition of network into parental species trees. The hybrid node is depicted as the circular point located on the right side of the arrow. The introgression scenario depicted on the left contains 2 parental species trees *S* and *I* at the middle and right. The probabilities assigned to the 2 parental species trees are $\gamma$ and $1-\gamma$.

Table S2. Parsimony-informative site patterns probabilities for outflow scenario.

| | ***BBAA*** | ***BABA*** | ***ABBA*** |
|---|---|---|---|
| 1 | $\frac{3}{64}$ | $\frac{3}{64}$ | $\frac{3}{64}$ |
| $e^{-\frac{8\tau_g}{3}}$ | $-\frac{3k'^{-1}\gamma}{64(1+\lambda a)}$ | $\frac{9k'^{-1}\gamma}{64(1+\lambda a)}$ | $-\frac{3k'^{-1}\gamma}{64(1+\lambda a)}$ |
| $e^{-\frac{8\tau_1}{3}}$ | $\frac{3k(3-\gamma k')}{64(1+a)}$ | $-\frac{3k(1+\gamma k')}{64(1+a)}$ | $\frac{3k(3\gamma k'-1)}{64(1+a)}$ |
| $e^{-\frac{8\tau_2}{3}}$ | $-\frac{3(1+k)(1-\gamma)}{64(1+a)}$ | $\frac{3(3k-1)(1-\gamma)}{64(1+a)}$ | $\frac{3(3-k)(1-\gamma)}{64(1+a)}$ |
| $e^{-\frac{8\tau_3}{3}}$ | $\frac{3(2-k)+9\gamma(kk'-1)}{64(1+a)}$ | $\frac{3(-kk'\gamma+\gamma-k+2)}{64(1+a)}$ | $\frac{3(-kk'\gamma+\gamma+3k-2)}{64(1+a)}$ |

| | | | |
|---|---|---|---|
| $e^{-\frac{4}{3}(\tau_g+2\tau_1)}$ | $-\frac{3k\gamma}{16(1+a)(2+\lambda a)}$ | $-\frac{3k\gamma}{16(1+a)(2+\lambda a)}$ | $-\frac{3k\gamma}{16(1+a)(2+\lambda a)}$ |
| $e^{-\frac{4}{3}(\tau_g+2\tau_3)}$ | $-\frac{3k\gamma}{16(1+a)(2+\lambda a)}$ | $-\frac{3k\gamma}{16(1+a)(2+\lambda a)}$ | $-\frac{3k\gamma}{16(1+a)(2+\lambda a)}$ |
| $e^{-\frac{4}{3}(\tau_1+2\tau_2)}$ | $-\frac{3k(1-\gamma)}{16(1+a)(2+a)}$ | $-\frac{3k(1-\gamma)}{16(1+a)(2+a)}$ | $-\frac{3k(1-\gamma)}{16(1+a)(2+a)}$ |
| $e^{-\frac{4}{3}(\tau_1+2\tau_3)}$ | $-\frac{3k(1+k'\gamma)}{16(1+a)(2+a)}$ | $-\frac{3k(1+k'\gamma)}{16(1+a)(2+a)}$ | $-\frac{3k(1+k'\gamma)}{16(1+a)(2+a)}$ |
| $e^{-\frac{4}{3}(\tau_2+2\tau_3)}$ | $-\frac{3(1+k)(1-\gamma)}{16(1+a)(2+a)}$ | $-\frac{3(1+k)(1-\gamma)}{16(1+a)(2+a)}$ | $-\frac{3(1+k)(1-\gamma)}{16(1+a)(2+a)}$ |
| $e^{-\frac{4}{3}(2\tau_g+2\tau_3)}$ | $\frac{3k'^{-1}\gamma}{64(1+a)(1+\lambda a)}$ | $\frac{27k'^{-1}\gamma}{64(1+a)(1+\lambda a)}$ | $\frac{3k'^{-1}\gamma}{64(1+a)(1+\lambda a)}$ |
| $e^{-\frac{4}{3}(2\tau_1+2\tau_3)}$ | $\frac{27k(1-\gamma)}{64(1+a)^2}$ | $\frac{3k(1-\gamma)}{64(1+a)^2}$ | $\frac{3k(1-\gamma)}{64(1+a)^2}$ |
| $e^{-\frac{4}{3}(\tau_g+\tau_1+2\tau_3)}$ | $\frac{3k\gamma}{4(1+a)(2+a)(2+\lambda a)}$ | $-\frac{3k\gamma}{4(1+a)(2+a)(2+\lambda a)}$ | $\frac{3k\gamma}{4(1+a)(2+a)(2+\lambda a)}$ |
| $e^{-\frac{4}{3}(\tau_1+\tau_2+2\tau_3)}$ | $-\frac{3k(1-\gamma)}{4(1+a)(2+a)^2}$ | $\frac{3k(1-\gamma)}{4(1+a)(2+a)^2}$ | $\frac{3k(1-\gamma)}{4(1+a)(2+a)^2}$ |
| $e^{-\frac{2(\tau_1-\tau_g)}{\theta}}e^{-\frac{8\tau_1}{3}}$ | $-\frac{3(\lambda-1)ak^2k'\gamma}{64(1+a)(1+\lambda a)}$ | $\frac{9(\lambda-1)ak^2k'\gamma}{64(1+a)(1+\lambda a)}$ | $-\frac{3(\lambda-1)ak^2k'\gamma}{64(1+a)(1+\lambda a)}$ |
| $e^{-\frac{2(\tau_1-\tau_g)}{\theta}}e^{-\frac{12\tau_1}{3}}$ | $-\frac{3(\lambda-1)ak^2k'\gamma}{16(1+a)(2+a)(2+\lambda a)}$ | $-\frac{3(\lambda-1)ak^2k'\gamma}{16(1+a)(2+a)(2+\lambda a)}$ | $-\frac{3(\lambda-1)ak^2k'\gamma}{16(1+a)(2+a)(2+\lambda a)}$ |
| $e^{-\frac{2(\tau_1-\tau_g)}{\theta}}e^{-\frac{4(\tau_1+2\tau_3)}{3}}$ | $-\frac{3(\lambda-1)ak^2k'\gamma}{16(1+a)(2+a)(2+\lambda a)}$ | $-\frac{3(\lambda-1)ak^2k'\gamma}{16(1+a)(2+a)(2+\lambda a)}$ | $-\frac{3(\lambda-1)ak^2k'\gamma}{16(1+a)(2+a)(2+\lambda a)}$ |
| $e^{-\frac{2(\tau_2-\tau_1)}{\theta}-\frac{4}{3}(2\tau_2+2\tau_3)}$ | $\frac{3ak(4+a)^2(1-\gamma)}{32(1+a)^2(2+a)^2(3+a)}$ | $\frac{3ak(32+40a+10a^2)(1-\gamma)}{64(1+a)^2(2+a)^2(3+a)}$ | $\frac{3ak(32+40a+10a^2)(1-\gamma)}{64(1+a)^2(2+a)^2(3+a)}$ |
| $e^{-\frac{2(\tau_1-\tau_g)}{\theta}}e^{-\frac{4}{3}(2\tau_1+2\tau_3)}$ | $3ak^2k'\gamma(-2(2+a)$ $+\lambda^2a(30+11a)$ $+\lambda(36+12a-a^2))$ $/64(1+a)(2+a)(3$ $+a)(1+\lambda a)(2+\lambda a)$ | $3ak^2k'\gamma(-2(26+9a)$ $+\lambda^2a(30+11a)$ $+\lambda(84+4a-9a^2))$ $/64(1+a)(2+a)(3+a)(1$ $+\lambda a)(2+\lambda a)$ | $3ak^2k'\gamma(-2(2+a)$ $+\lambda^2a(30+11a)$ $+\lambda(36+12a-a^2))$ $/64(1+a)(2+a)(3+a)(1$ $+\lambda a)(2+\lambda a)$ |

Note: $k=e^{-(4/3)(\lambda-1)\tau_1}, k'=e^{-(4/3)(1-\lambda)\tau_g}$, a $=4\theta/3$.

**Supplementary Note 3**

When $\gamma = 0$, there is:

$$P(BBAA) - P(BABA) = \frac{3}{64}(\frac{4k}{(1+a)}(e^{-\frac{8\tau_1}{3}} - e^{-\frac{8\tau_2}{3}}) + \frac{8k}{(1+a)^2}e^{-\frac{4}{3}(2\tau_1+2\tau_3)} - \frac{32k}{(1+a)(2+a)^2}e^{-\frac{4}{3}(\tau_1+\tau_2+2\tau_3)} - \frac{8ka^2}{(1+a)^2(2+a)^2}e^{-\frac{2(\tau_2-\tau_1)}{\theta}-\frac{4}{3}(2\tau_2+2\tau_3)})$$

$$\because \frac{4k}{(1+a)}(e^{-\frac{8\tau_1}{3}} - e^{-\frac{8\tau_2}{3}}) > 0 \; and$$

$$\frac{8k}{(1+a)^2}e^{-\frac{4}{3}(2\tau_1+2\tau_3)} - \frac{32k}{(1+a)(2+a)^2}e^{-\frac{4}{3}(\tau_1+\tau_2+2\tau_3)} > (\frac{8k}{(1+a)^2} - \frac{32k}{(1+a)(2+a)^2})e^{-\frac{4}{3}(\tau_1+\tau_2+2\tau_3)} = \frac{8ka^2}{(1+a)^2(2+a)^2}e^{-\frac{4}{3}(\tau_1+\tau_2+2\tau_3)}$$

$$\therefore P(BBAA) - P(BABA) > \frac{3}{64}(\frac{8ka^2}{(1+a)^2(2+a)^2}e^{-\frac{4}{3}(\tau_1+\tau_2+2\tau_3)} - \frac{8ka^2}{(1+a)^2(2+a)^2}e^{-\frac{2(\tau_2-\tau_1)}{\theta}-\frac{4}{3}(2\tau_2+2\tau_3)}) > 0$$

### Supplementary Note 4

In cases of $\gamma = 0$, when $\tau_1, \tau_2 \to 0$, and $\lambda\tau_1, \tau_3 \to \infty$, there is:

$$P(BABA) \to \frac{3}{64}(1 - \frac{1}{1 + 4/3\theta})$$

$$P(ABBA) \to \frac{3}{64}(1 + \frac{3}{1 + 4/3\theta})$$

$$\Rightarrow D = \frac{P(ABBA) - P(BABA)}{P(ABBA) + P(BABA)} \to \frac{4}{1 + 4/3\theta}/(2 + \frac{2}{1 + 4/3\theta})$$

$$if\ \theta \ll 1, D \to 1.$$

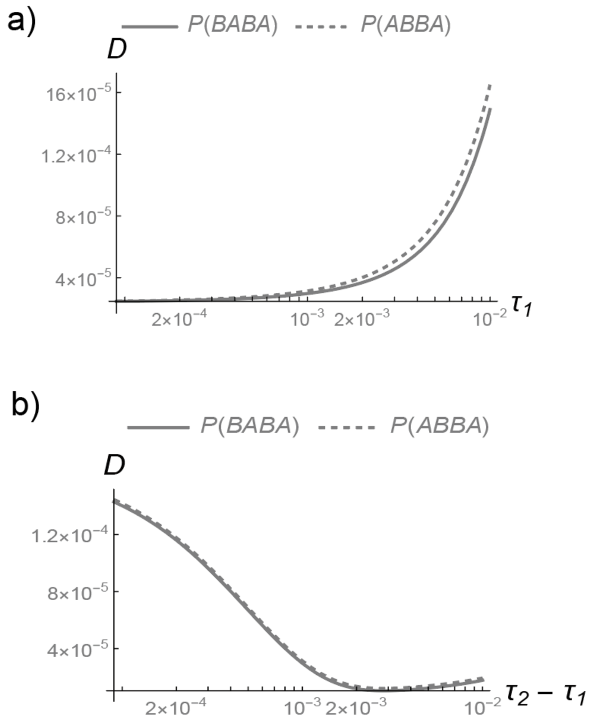


Figure S3. The frequencies of *ABBA* and *BABA* site patterns as $\tau_1$ and $\tau_2 - \tau_1$ vary.

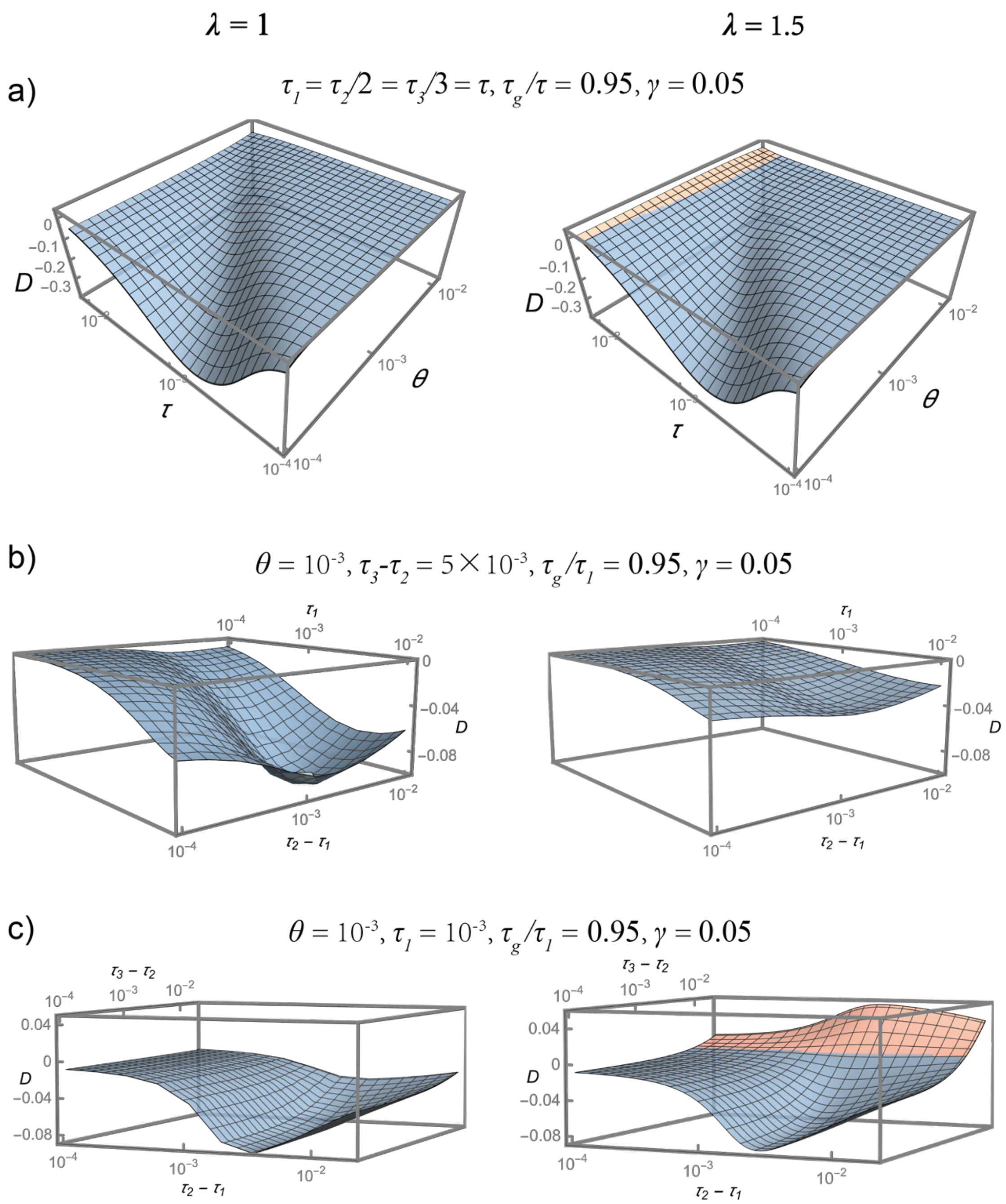


Figure S4. The effects of $\theta$ and the branch lengths $\tau_1$, $\tau_2-\tau_1$ and $\tau_3-\tau_2$ on *D*-values in the presence of introgression.

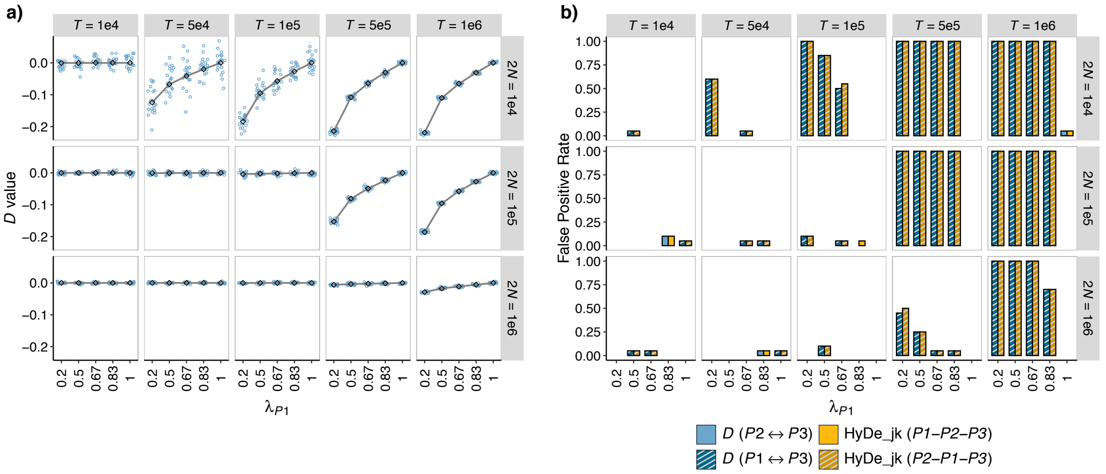


Fig S5. Interactive effects of phylogenetic depth ($T$), population size ($2N$), and rate variation on introgression testing. The simulated scenarios correspond to Figure 3a. The values on the strips at the top and right of each plot indicate the phylogenetic depth ($T$) and population size ($2N$), respectively. $\lambda_{P1}$ is labeled on the x-axis. a) $D$-values: Colored points represent $D$-value estimates, with solid lines indicating the theoretical expected $D$-values. b) False-positive rate of $D$-statistic and HyDe_jk.

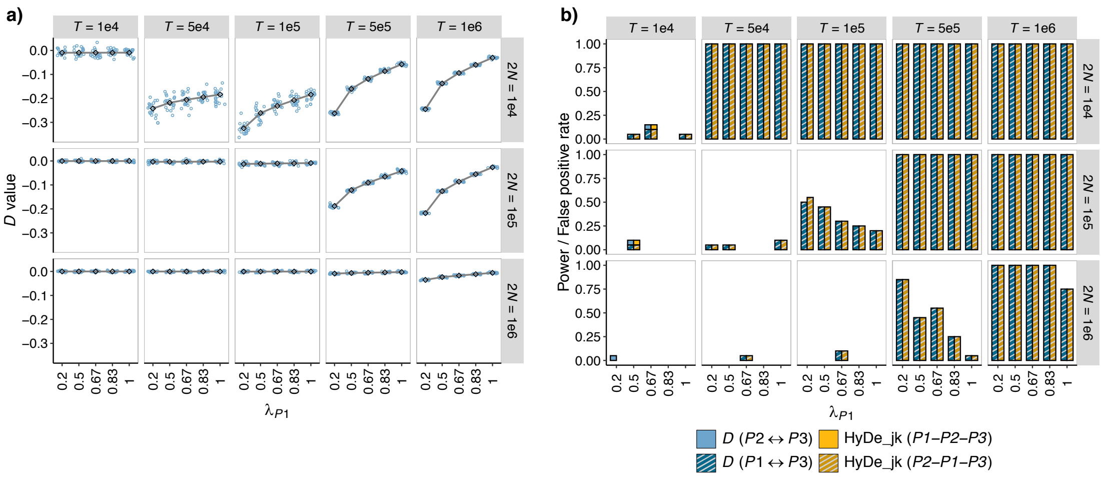


Fig S6. Effect of rate variation on performance of *D*-statistic and HyDe, in presence of introgression. The simulated scenarios correspond to Figure 7a. The values on the strips at the top and right of each plot indicate the phylogenetic depth (*T*) and population size (2*N*), respectively. $\lambda_{P1}$ is labeled on the x-axis. a) *D*-values: Colored points represent *D*-value estimates, with solid lines indicating the theoretical expected *D*-values. b) False-positive rate of *D*-statistic and HyDe_jk.

### Reference


Gardner EM, Bruun-Lund S, Niissalo M, Chantarasuwan B, Clement WL, Geri C, Harrison RD, Hipp AL, Holvoet M, Khew G. 2023. Echoes of ancient introgression punctuate stable genomic lineages in the evolution of figs. Proc. Natl. Acad. Sci. U.S.A. 120:e2222035120.

Graur D, Li W-HL. 2000. Fundamentals of molecular evolution. 12th ed. Sunderland, MA: Sinauer.

Karimi N, Grover CE, Gallagher JP, Wendel JF, Ané C, Baum DA. 2020. Reticulate evolution helps explain apparent homoplasy in floral biology and pollination in baobabs (*Adansonia*; Bombacoideae; Malvaceae). Syst. Biol. 69:462–478.

Leduc-Robert G, Maddison WP. 2018. Phylogeny with introgression in *Habronattus* jumping spiders (Araneae: Salticidae). BMC Evol. Biol. 18:1–23.

Liu BB, Ren C, Kwak M, Hodel RG, Xu C, He J, Zhou WB, Huang CH, Ma H, Qian GZ. 2022. Phylogenomic conflict analyses in the apple genus *Malus* sl reveal widespread hybridization and allopolyploidy driving diversification, with insights into the complex biogeographic history in the Northern Hemisphere. J. Integr. Plant Biol. 64:1020–1043.

Meng C, Kubatko LS. 2009. Detecting hybrid speciation in the presence of incomplete lineage sorting using gene tree incongruence: A model. Theor. Popul. Biol. 75:35–45.

Richards A, Kubatko L. 2022. Site pattern probabilities under the multispecies coalescent and a relaxed molecular clock: Theory and applications. J. Theor. Biol. 542:111078.

Sarver BA, Herrera ND, Sneddon D, Hunter SS, Settles ML, Kronenberg Z, Demboski JR, Good JM, Sullivan J. 2021. Diversification, introgression, and rampant cytonuclear discordance in rocky mountains chipmunks (Sciuridae: *Tamias*). Syst. Biol. 70:908–921.

Wu M, Kostyun JL, Hahn MW, Moyle LC. 2018. Dissecting the basis of novel trait evolution in a radiation with widespread phylogenetic discordance. Mol. Ecol. 27:3301–3316.

Zhu S, Degnan JH. 2017. Displayed trees do not determine distinguishability under the network multispecies coalescent. Syst. Biol. 66:283–298.
